# Tissue damage-induced axon injury-associated responses in sensory neurons: requirements, prevention, and potential role in persistent post-surgical pain

**DOI:** 10.1101/2025.02.11.637320

**Authors:** Kristofer K. Rau, Benjamin J. Harrison, Gayathri Venkat, Sara E. Petruska, Bradley K. Taylor, Caitlin E. Hill, Jeffrey C. Petruska

## Abstract

Pain resulting from tissue damage, including surgical incision, is often only partially responsive to standard treatments focusing on inflammation, suggesting additional mechanisms are involved. Tissue damage leads to expression in dorsal root ganglion (DRG) sensory neurons of activating transcription factor 3 (ATF3), a known injury-induced transcription factor. ATF3 expression is associated with sensitization of cellular physiology and enhanced amplitude/duration of a nociceptive reflex. It is unclear how tissue damage leads to these changes in the sensory neurons, but it could include direct damage to the tissue-innervating axons and inflammation-associated retrograde biochemical signalling. We examined the necessity and sufficiency of incision, inflammation, and axonal conduction for induction of ATF3 in response to skin incision. Incision outside of a single dermatome, but close enough to induce inflammation inside the dermatome, was not sufficient to induce ATF3 expression in the corresponding DRG. Incision inside the dermatomeled to strong expression of ATF3. Anti-inflammatory treatments did not prevent this induction of ATF3. In rodent models of repeated injury - a major etiological factor for chronic pain - ATF3 expression was synergistically increased and the threshold for paw-withdrawal to mechanical stimulation was significantly decreased for an extended duration. Together, these results suggest that actual damage to axons innervating the skin is both necessary and sufficient for induction of ATF3, expression of which appears additionally increased by repeated injury. Further, pre-treatment of the nerves innervating the incised skin with bupivacaine, a local anesthetic commonly used to reduce surgical pain, did not prevent induction of ATF3, indicating that conduction of action potentials is not necessary for induction of ATF3. We also determined that closure of incision with surgical glue significantly reduced incision-induced expression of ATF3 and GAP-43. Intriguingly, treatment with polyethylene glycol (PEG), known to enhance membrane integrity after injury among other effects, reduced incision-associated ATF3 expression and electrophysiological changes. These results suggest that pain resulting from tissue damage may arise from a mix of ATF3-independent inflammation-related mechanisms as well as ATF3-/axonal-damage-associated mechanism and therefore require a mix of approaches to achieve more complete control some of which we suggest here..

**Funding:** JCP NIH R01NS109936, R21NS120498, KSCHIRT 10-10 BJH NIH R01NS121533

**SCOPE statement:** Pain resulting from tissue damage, including surgical incision, is often only partially responsive to anti-inflammatory treatments, suggesting multiple mechanisms at work, including neuropathic. Cutaneous tissue damage leads to expression in DRG sensory neurons of the injury marker ATF3 - associated with physiological sensitization and enhancement of a nociceptive reflex. We examined some of the conditions for induction of ATF3 in response to incision of skin and the impact of interventions. Treatment with anti-inflammatory ketoprofen and/or local anesthetic did not prevent the induction of ATF3, together suggesting that actual damage to axons innervating the skin is both necessary and sufficient for induction of ATF3. Repeated incision induced unique changes in expression of ATF3- and pain-associated genes. Closure with surgical glue reduced incision-induced gene expression compared to closure with staples. Treatment with polyethylene glycol (PEG), known to enhance membrane integrity after axonal injury, reduced induction of ATF3 and electrophysiological changes. These experiments were designed to identify distinct pain-related mechanisms with pre-clinical animal models that reflect existing clinical practice and feasible future practice. These results suggest that pain resulting from tissue damage likely arises from mixed mechanisms – including neuropathic – and therefore require a mix of approaches to achieve more complete control.

## Introduction

Pain associated with tissue damage, including surgery, is generally treated as inflammatory pain (e.g., [1–3]). There is certainly a significant component of the pain, especially in the acute and sub-acute setting, that is due to inflammation-induced sensitization of sensory neurons. However, in many cases the pain is only partially controlled by anti-inflammatory drugs and/or outlasts healing of the wound and resolution of inflammation. Etiologically, the most documentable and measurable form of tissue damage-related pain is persistent post-surgical pain (PPP). PPP does not occur in all patients, but one study identified that moderate-severe pain occurs in 15-30% of patients more than 2 years after surgery [4], and a recent review suggests 5-85% [5], with rates varying with different types of surgeries. Although it is possible that we will arrive at some new understanding of the known mechanisms that will account for these occurrences, it is more likely that additional currently-unappreciated mechanisms will provide a more thorough explanation.

The field acknowledges a neuropathic component to most persistent post-surgical pain, which in common practice refers to injury to nerves. Certainly there are many cases of persistent post-surgical pain in which overt nerve damage occurred, but this is not always the case. It is possible that there was covert nerve damage in those cases, but it is also possible that there is another source of damage that leads to a response in the sensory neurons that generates a nerve injury-like response without injury to a discrete nerve.

Mechanistically, it is recognized that persistent pain “…is neither the result of an inflammatory process alone nor only the result of isolated injury to nerves” [6]. This suggests that there are additional non-inflammation-related mechanisms at work, potentially including neuropathic mechanisms. Despite a recognition of a clear role of role of various forms of nerve injury in the etiology of persistent post-surgical pain, the field appears apprehensive to expand this factor to recognize *axonal injury* as a contributing factor, evidenced by its absence in otherwise comprehensive reviews (e.g., [5, 7]). Revising the contributing factor to be axonal injury, as opposed to nerve injury, would recognize the role of injury to sensory (and possibly autonomic and motor) neurons yet still encompass both injury to nerve tissue and peripheral target tissue.

The identification of nerve injury-like molecular and functional responses associated with tissue damage can offer new insights into the underlying causes of clinical pain, but it also raises many questions regarding necessary and sufficient conditions, particularly regarding common clinical practice. Further, if these responses are part of the overall pain experience, can they be prevented, reversed, or treated similarly to many inflammatory mechanisms, or are different treatments/preventions necessary?

Tissue damage can induce long-lasting changes in gene expression and physiology in sensory neurons that appear very similar to those induced by nerve injury – a known cause of neuropathic pain. This was reported in models of skin incision [8–10], joint degeneration [11–13], dry eye [14], and therapeutic radiation treatment [15]. We therefore sought to examine what clinically-modifiable factors might influence the expression of these injury-like responses in sensory neurons. We examined the influence of distance of injury from innervation zone, local anesthetics, anti-inflammatories, and closure methods on the expression of ATF3. Because repeated injury is often considered a risk factor for developing persistent pain, we also examined the impact of repeated injury on nociceptive behaviors and expression of ATF3 and relevant pain-related genes.

Working from the principle that tissue damage can induce axolemma disruption, we also examined the effects of the “fusogenic” agent polyethylene glycol (PEG). PEG undoubtedly has many effects, but it is clear that it can induce sealing of damaged membranes, even to the extent of – at least temporarily – re-annealing/fusing the cut ends of axons, restoring conduction after nerve transection (e.g., [16–20]) and enhancing functional outcomes after spinal cord injury (e.g., [21–25]).

## METHODS

### Animals

Animal care and procedures were carried out at the University of Louisville and the University of Kentucky and were in accord with approved IACUC protocols. Age-matched adult female Sprague-Dawley rats (180–200 g; Taconic, Indianapolis, Indiana) and age-matched male and female C57BL-6 mice (Charles River) were used for these studies [Real-time quantitative PCR (RT-qPCR) and electrophysiology]. For all surgeries using rats, animals were anesthetized with ketamine/xylazine (i.p. injection; 80 mg/kg ketamine; 10 mg/kg xylazine) and body temperature was monitored and maintained at 36 °C throughout the surgeries which lasted 20–30 min. Puralube ointment (Dechra) was used to protect the rat’s eyes during surgery. Following surgery, rats were given lactated ringer’s solution (5 ml, i.p.) to prevent dehydration, and gentamycin (Gentafuse; 0.1 ml, i.m., every other day for 7 days) to prevent infection. Rats ages 4-5 weeks at the beginning of experiments were housed individually throughout the experiment. Mice ages 6–16 weeks at the beginning of experiments were housed 2–4 per cage. All animals were monitored daily, maintained on a 12 h light/dark cycle at 20–22°C and 45 ± 10% relative humidity, with food and water provided ad libitum.

### Surgical procedures

Incisions involved cutting the full thickness of both the hairy skin and the underlying attached cutaneous trunci muscle. Following incision, the skin was closed with Ethilon nylon suture (5-0, Ethicon), and coated with Bacitracin antibiotic ointment (Actavis) to prevent infection. The experimental incision used to examine the molecular changes in the DRG neurons was made on the left side of the rats. It was located 1 cm lateral to the vertebral column and extended parallel to the vertebral column for 3 cm (including approximately the T7–12 dermatomes). The location of the incision ensured that the dorsal cutaneous nerves were not damaged by the incision. In experiments utilizing polyethylene glycol (PEG; 30% w/w in Ringer’s solution; Sigma), 1 cc of PEG was applied post surgically to the surface of the wound site. An additional 1 cc of PEG was injected IP.

### Tissue collection

The range of time points examined (4–28 days post-incision) encompasses both sub-acute time points in which inflammation-related mechanisms occur, and later time points when inflammation has typically resolved (≥ 10–14 days). This range also encompasses the majority of the wound healing process. The skin is closed (surface wound contraction) to the point where sutures and/or staples can be removed by 7–10 days. Rats received an experimental incision and tissue was collected 4, 7, 14 or 28 days post-incision (DPI: per time point). A control group of rats received no experimental incision. For tissue collection, rats were euthanized with an overdose of pentobarbital and transcardially exsanguinated with heparinized phosphate buffered saline (PBS; pH 7.4). This was followed by 33% vol/vol RNAlater (Qiagen) in heparinized PBS to help preserve the RNA. Three adjacent DRGs with projections to the incision site (typically T10–T12; innervation of incision site confirmed by gross-anatomical dissection) were collected, pooled together, and placed in 100% RNAlater overnight at 4 °C and then stored at −80 °C until RNA isolation.

### RNA isolation

To isolate RNA from the DRGs, samples were placed on ice and 350 μl RLT lysis buffer (Qiagen) and 2-mercaptoethanol was added. Tissue was homogenized for 1 min using a motorized dual Teflon glass homogenizer (Kontes). RNA was extracted using the RNeasy plus micro kit (Qiagen) as per manufacturer’s protocol. Genomic DNA was removed using the DNA eliminator affinity spin column and RNA was purified by affinity purification using RNA spin columns. Samples were eluted in 14 μl of nuclease free water. RNA integrity was assessed by UV spectrometry and the Bioanalyzer (Agilent Technologies). RNA samples with 260 nm/280 nm ratios above 1.9 and 260 nm/230 nm ratios and RNA integrity numbers above 1.8 were used for RT-qPCR.

### RT-qPCR

RT-qPCR was used to quantify the expression of genes within DRGs following skin incision. We examined ATF3 and GAP-43. cDNA was generated from the RNA samples using the Quantitect first strand synthesis kit (Qiagen) according to the manufacturer’s protocol. For each PCR reaction, 5 ng of cDNA template was used. Samples were run in triplicate, and control reactions (without template) were included with every amplification run. SYBR green RT-qPCR was carried out using a Rotorgene real time PCR detection instrument (Corbett Research). Relative fold-changes of RNA were calculated by the ΔΔCT method using GAPDH as the stable internal reference gene. Small differences in RT-qPCR reaction efficiency between primer sets were accounted for using the standard curve quantification methods.

#### Gene Forward Primer (5’-3’) Tm (°C) Reverse Primer (5’-3’) Tm (°C)

GAPDH ATGGCCTTCCGTGTTCCTAC 65.0 AGACAACCTGGTCCTCAGTG 61.9

ATF3 GAGATGTCAGTCACCAAGTC 60.1 TTCTTCAGCTCCTCGATCTG 61.4

GAP43 CTAAACAAGCCGATGTGCC 63.2 TTCTTTACCCTCATCCTGTCG 62.9

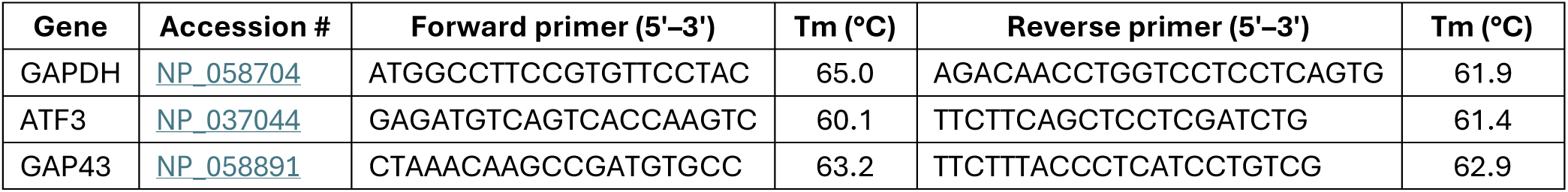

### Tracer incision

For rats in which DRGs were to be examined electrophysiologically, an additional skin incision was made, on the right side, seven days prior to the experimental skin incision to allow for tracer injection and transport. This skin incision to enable tracer injection was placed on the right side to prevent injury to the axons of interest on the left side. Following incision on the right side, the skin was reflected to expose the underside of the contralateral (left) skin. Depending on the experimental objective, 0.5% DiI (1,1′-dilinoleyl-3,3,3′,3′-tetramethylindocarbocyanine perchlorate; 5 mg FastDiI dissolved in 1 ml methanol; Invitrogen) was injected into either CTM or the subdermal layer of the skin using a Hamilton syringe. Ten injections of 1 μl each, were used to target the terminal field as described previously [10]. The Experimental Incision was approximately 5 mm distal to, and extended at least two dermatomes rostral and caudal from the DiI injected region. Preliminary studies indicated that this method maximized the percentage of DiI-labeled cells that were injured by incision, as indicated by ATF3/DiI co-localization.

### DRG dissociations for electrophysiology

DRGs were isolated, dissociated, and plated following the method described by Petruska et al. (2000, 2002). Enzymatic digestion of the DRGs was performed using dispase (neutral protease, 5 mg/ml; Boehringer Mannheim) and collagenase (type 1, 2 mg/ml; Sigma) in Tyrode’s solution for 90 minutes at 35 °C. To facilitate dissociation, the DRGs were gently triturated every 30 minutes. After enzymatic digestion, the cells were centrifuged at 800 rpm for 3 minutes, resuspended in fresh Tyrode’s solution, and then plated onto poly-L-lysine (Sigma)-coated dishes. The dishes were kept in an aerated holding bath for at least 2 hours before recording. Recordings were conducted within 10 hours of DRG retrieval from the animal, a time frame that precedes the translation of ATF3 (Rau et al., 2014). Therefore, the observed ATF3 expression in dissociated/recorded neurons was attributed to the skin incision rather than the dissociation process.

### Electrophysiological recording

Whole-cell patch recording was used to determine the electrophysiological properties of individually isolated DiI-labeled neurons, specifically those projecting to the site of the skin incision. The electrophysiology procedures followed previously detailed methods (Petruska et al., 2000, 2002) and were conducted using a Scientifica SliceScope Pro system. Electrodes (2–4 MΩ) were prepared from glass pipettes using a horizontal puller (Sutter model P1000).

The extracellular solution (Tyrode’s) consisted of 140 mM NaCl, 4 mM KCl, 2 mM MgCl2, 2 mM CaCl2, 10 mM glucose, and 10 mM HEPES, with the pH adjusted to 7.4 using NaOH. The recording electrodes were filled with a solution of 120 mM KCl, 5 mM Na2-ATP, 0.4 mM Na2-GTP, 5 mM EGTA, 2.25 mM CaCl2, 5 mM MgCl2, and 20 mM HEPES, also adjusted to pH 7.4 with KOH and with an osmolarity of approximately 315– 325 mOsm.

DiI-labeled neurons were identified using brief illumination with epifluorescence microscopy (total exposure of field <1 min). Once identified, whole-cell recordings were conducted using an Axoclamp 2B (Molecular Devices). Stimuli were controlled, and digital records were captured using pClamp10 software and a Digidata 1440 (Molecular Devices). Series resistance (RS) was compensated by 50–70%. Whole-cell resistance (RM) and capacitance were assessed using voltage transients associated with small step commands (10 mV) via pClamp software. All experiments were carried out at room temperature, and only cells with a resting membrane potential (RMP) of −40 to −70 mV were included.

After obtaining and stabilizing individual DRG neurons in whole-cell patch-clamp voltage-clamp mode, the cells were switched to current-clamp mode to assess changes in membrane potential. To evaluate cellular excitability, rheobase and action potential frequency in response to standardized depolarizing current steps were acquired. Action potentials were evoked with a 1 ms, 2 nA current step, and the average of ten action potentials was used to determine the after-hyperpolarization duration (80% recovery to baseline) and amplitude (voltage decrease from RMP to the lowest point of the after-hyperpolarization) as per Djouhri et al. (1998). Action potential threshold and duration at threshold (APDt) were measured at the rheobase using 500-ms square pulses, increased in 50-pA increments every 2 seconds and refined further with 5-pA increments. APDt was measured from the first upward deflection of the action potential waveform to its return to the threshold membrane potential. The number of evoked action potentials and peak instantaneous frequency were determined using increasing voltage steps (1-second stimulus duration, 10-second interstimulus interval; stimuli increased in 50-pA steps over 20 sweeps, resulting in evoked current recordings from 50 to 1000 pA). Electrophysiological data were analyzed using Clampfit software.

Only one cell was recorded per dish. After recording, the cell’s location was marked by physically noting it on the underside of the plastic culture dish and capturing a digital image using a Scientifica monochrome camera and SCIght 2.0 software. The bath solution was then replaced with 4% PFA in PBS for 10 minutes, followed by rinsing and replacing with 100% PBS solution. Cells were stored at 4 °C until immunolabeling procedures were carried out. Immunocytochemistry was performed to examine ATF3 expression in recorded DRG neurons.

### Mouse plantar incision

Animals were housed in a temperature-controlled room on a 14/10 h light/dark cycle and were provided food and water ad libitum. Mice were acclimated to the colony housing room for at least 4 d and then acclimated to handling for 3 min per day on each of 4 d before the initiation of the experiments. All animal-use protocols were approved by the Institutional Animal Care and Use Committee at the University of Kentucky.

Surgical procedures were as previously described [26, 27]. Briefly, postoperative hyperalgesia was induced by longitudinal incision of the plantaris muscle. Anesthesia was by isoflurane (5% induction and 1.5–2% maintenance via a nose cone) and antisepsis by Chlorascrub then alcohol to the left hindpaw. A no. 11 scalpel blade was used to make a 5 mm long incision through the skin and fascia, beginning 2 mm proximal from the end of the heel and extending distally toward the digits. The underlying muscle was raised with curved forceps and then incised longitudinally, leaving the terminal connective tissue intact. The overlying skin was closed with synthetic 5–0 sutures (PDS* II, Ethicon), followed by application of antibiotic ointment. Sutures were removed on postoperative day 10. Sham controls received anesthesia but no surgical incision.

### Mouse withdrawal reflex

Animals were acclimated to a temperature- and light-controlled room within individual Plexiglas bottomless boxes placed on the top of a stainless steel mesh platform for 30–60 min before behavioral testing. Mechanical thresholds were assessed using an incremental series of eight von Frey filaments (Stoelting) of logarithmic stiffness (0.008–6 g). The 50% withdrawal threshold was determined using an up-down method [28]. Each filament was applied perpendicular to the surface of the skin just lateral to the incision site with enough force to cause a slight bending of the filament. A positive response was defined by a quick withdrawal of the paw within 5 s. Gram force was logarithmically converted to 50% mechanical threshold. Thresholds were measured from baseline through 28 d after the first plantar incision model (PIM) to confirm the development and resolution of mechanical allodynia and then again after the second incision and for 21 days thereafter in the 2-incision group. Control groups had corresponding single incision or no-incision.

### Bioinformatics

Putative ATF3-regulated human genes were identified from the GSEA website (https://www.gsea-msigdb.org/) using data from TRANSFAC (v7.4). These were cross-referenced against genes assigned with ontology annotations (from AmiGO): “regulation of membrane potential” (biological process, GO:0042391), “response to pain” (biological process, GO:0048266), and/or “sensory perception of pain” (biological process, GO:0019233).

### Statistical analyses

Statistical analyses were performed using SigmaPlot/SigmaStat (Systat Software, San Jose, CA, USA). First-pass analysis to examine differences between the skin-incised and control groups for all assessments was done using analysis of variance (ANOVA) or repeated measures ANOVA and was followed by pair-wise comparisons (Student-Newman-Keuls). Differences were considered to be statistically significant if p < 0.05. Data is presented as mean ± standard deviation.

### Terminology

In this report we use terms that are often considered interchangeable, but we make distinctions which we believe are, or will be, meaningful.

Injury vs. Damage: We use “injury” to imply the action, and “damage” to imply the result/condition.

Tissue damage vs. Incision: We are considering “incision” as one means, among many, of inducing “tissue damage”. “Incision” refers to clinical surgical practice and the current model and data, while “tissue damage” refers to the broader meaning and impact.

Nerve injury and axon injury: We consider these from the perspective of Gross Anatomy and Histology, where these are considered related but distinct due to different features, predominantly of the cellular complement. “Nerve injury” refers to injury of the Gross Anatomical structure of a peripheral nerve, whereas “axon injury” refers to injury to axons irrespective of the tissue in which they are resident. Further details are provided in Discussion.

## RESULTS

### Nerve conduction in the acute phase of tissue damage is not required for ATF3 upregulation

Local and regional anesthesia is routine and indispensable for many surgical procedures and to offer temporary relief for painful conditions [29]. Although this approach largely provides excellent relief/prevention of pain in the short-term, preventing the emergence of longer-term pain has had mixed results [29]. If the induction of an injury-like/cell-stress-like response in sensory neurons – indicated by expression of ATF3 – is part of the mechanism of persistent pain, then one would expect that ATF3 expression might occur irrespective of the use of conduction-blocking drugs.

We sought to determine if clinically-modeled treatment with local anesthetics could prevent induction of ATF3. As indicated in **Figure 1** we performed a longitudinal incision to the right of midline, beyond the reach of any left-side cross-midline innervation. This initial incision serves both to expose the left-side dorsal cutaneous nerves for accurate peri-nerve administration of the anesthetics, but also served as the positive control incision affecting many segments on the right side of the animal. We then applied a single bolus of bupivacaine (0.5%, clinical formulation) to create a fascia-contained dome of fluid surrounding a 3-5mm portion of each of the T9, 10, 12, and 13 dorsal cutaneous nerves.

We then identified the regions of skin that were insensate by using light pinch or prick with the sharp tips of #5 forceps and observing for the Cutaneous Trunci Muscle Reflex (CTMR)[30–33]. The CTMR is resistant to the pentobarbital anesthesia used for surgical procedures with these animals. The regions of insensate skin were marked and incisions made inside those insensate regions (**Figure 1**). Incisions were made toward the edge of the insensate zones to avoid cutting the dorsal cutaneous nerve trunks themselves as they penetrated the skin.

To address the unlikely-but-possible scenario that this approach might block conduction by fluid compressing the nerve (and thus potentially inducing ATF3 expression on its own) we performed a similar application of saline around unused dorsal cutaneous nerves. The CTMR could still be driven by stimuli applied to skin regions innervated by those nerves, indicating continuity of signal conduction.

As expected, ATF3 expression in the right-side DRG samples was strongly upregulated. It was also clear that clinically-modelled use of local anesthetic did not prevent incision-induced expression of ATF3, as the expression in left-side DRG samples was also strongly upregulated.

It is possible that the anesthetic may have modulated the ATF3 response to some degree, but making such a comparison in this surgical model is not entirely appropriate as the incisions on the left and right side were different and likely affected different numbers of sensory neurons. The functional control we performed strongly indicated that the peri-nerve bolus application of anesthetic did not compress the nerve. This is further suggested by the similarity of ATF3 expression between the positive control (skin-incision) and the experimental (incision+anesthetic) conditions - if the injection had compressed the nerve the magnitude of ATF3 expression may have been dramatically greater.

**Figure 1.**
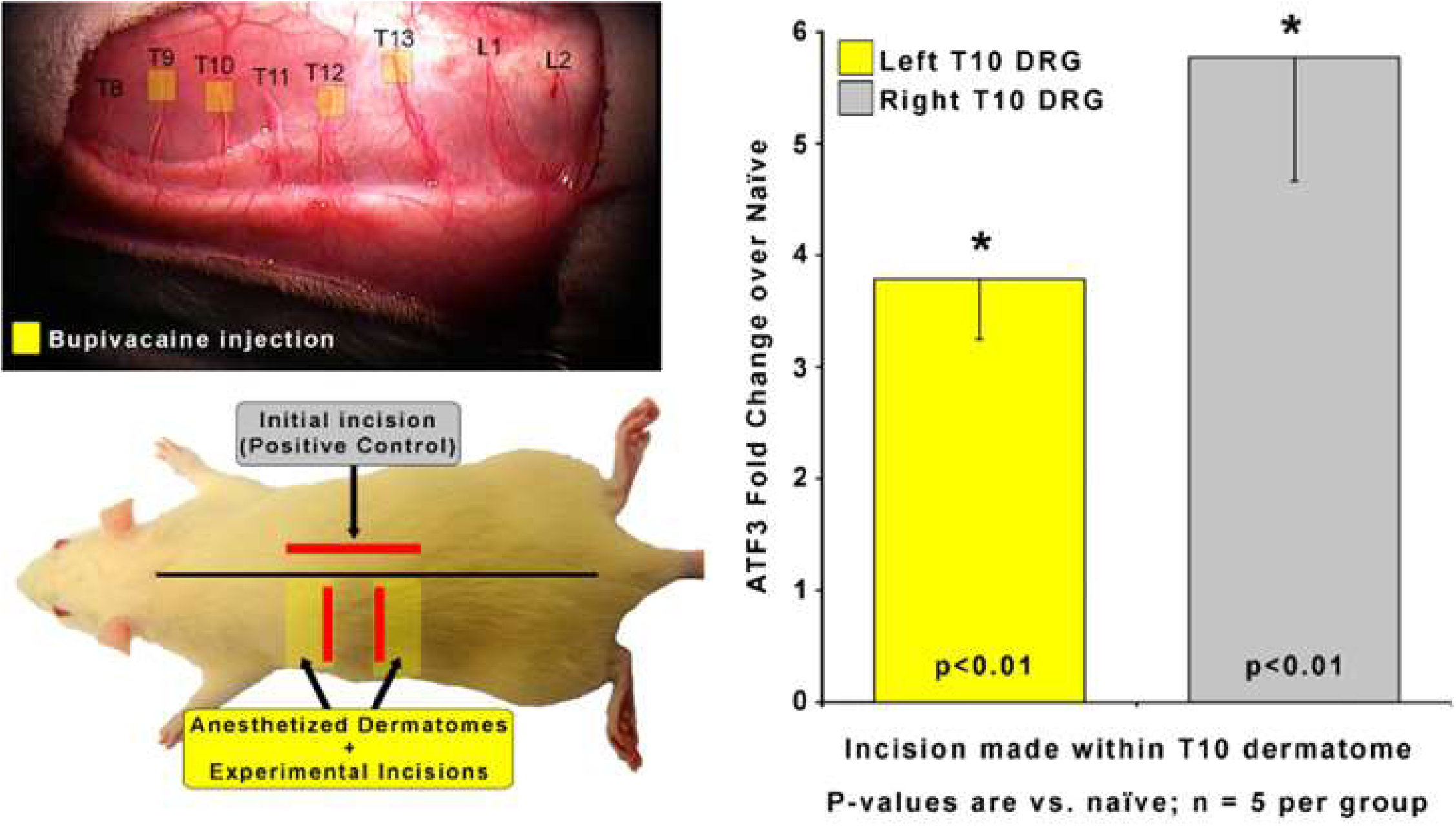
To determine if nerve conduction is required for ATF3 expression, a skin incision was made after injection of local anesthetic. ATF3 mRNA expression on the right. P-values provided are for post-hoc t-test vs. naive. **Clinical relevance:** Bupivacaine (as well as other anesthetics) is commonly used as a nerve block given prior to surgery to prevent nociception. Despite nerve block, ATF3 is still upregulated.

### Axonal damage is required for ATF3 upregulation

In order to assess the requirement for axonal injury, as opposed to just an inflammatory environment, we examined whether inflammation alone could induce ATF3 expression in DRG. We anesthetized the T9, 10, 12, 13 dorsal cutaneous nerves to allow us to define the T11 dorsal cutaneous nerve receptive field using the CTMR. Once spatially defined, we created a full-thickness incision that followed the (generally linear) border of the T11 dermatome. For these incisions, we borrowed from the findings of [34] which indicated that inflammation-mediating proteins were found in high concentration in the skin up to at least 1mm away from an incision, but were significantly lower (though still increased) at 2mm. Working from their data, our design included a group that should have had axons from the Left T11 DRG exposed to levels of inflammatory mediators varying from high (1mm) to low/nil (5mm). If inflammation alone were sufficient to induce ATF3 expression in sensory neurons, we would expect to see ATF3 expressed in the DRGs from animals with incisions 1mm outside the T11 dermatome but not from those with incisions 5mm outside the T11 dermatome. Instead, we found that none of the left-side DRGs expressed ATF3 above naive levels, indicating that presence of inflammatory mediators alone is not sufficient to induce expression of ATF3 (**Figure 2**). This indicates that actual damage of axons innervating the injured tissue is necessary for induction of ATF3 expression in the sensory neurons.

**Figure 2.**
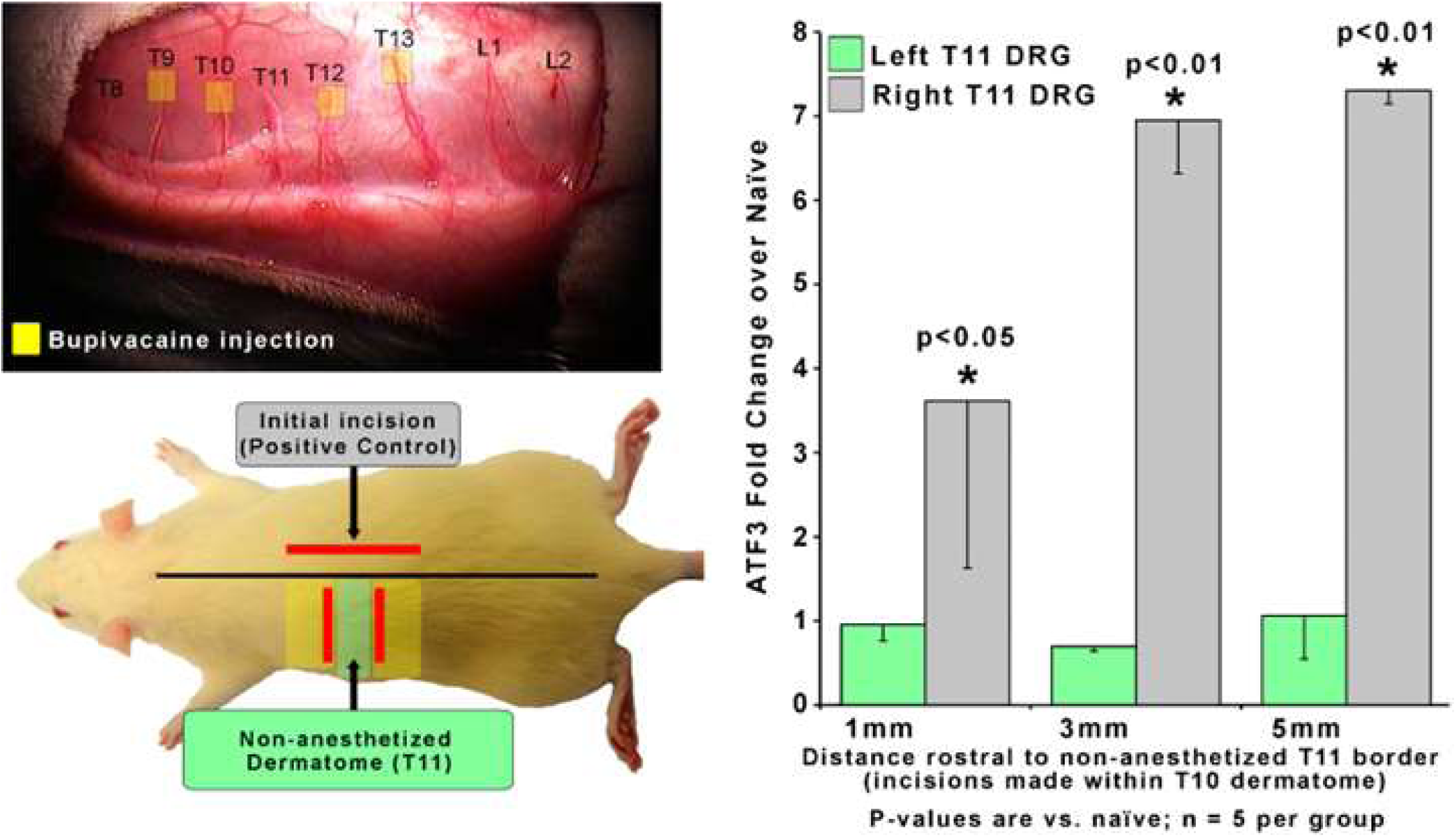
To determine if axonal damage is necessary for ATF3 expression, an incision was made 1, 3, or 5 mm outside of the non-anesthetized dermatome (T11). ATF3 mRNA expression is shown on the right. P-values are for post-hoc t-test vs. naive. **Clinical relevance:** Tissue reaction processes (including inflammation) in the region of overt tissue damage (as modeled here) are not sufficient to induce the injury/stress gene regulation response in sensory neurons innervating the tissue.

### Suppressing inflammation does not prevent ATF3 upregulation

It appears that inflammation is likely not sufficient to induce ATF3 expression, at least as induced here and at the time point we assessed. In order to determine if inflammation might still be necessary for tissue damage-induced ATF3 expression, we examined whether administration of an anti-inflammatory drug [35] might regulate ATF3 expression after incision. We further assessed whether a combination of anti-inflammatory drug and local anesthetic administration – a common clinical pain-control regimen – could prevent ATF3 expression.

The incisions for these experiments were longitudinal – parallel to the midline – so we included additional animals with similar treatments as reported in **Figure 1** to provide a more accurate comparison (Group 1). The induction of ATF3 expression in DRG innervating incised skin was not prevented or reduced by anti-inflammatory treatment (**Figure 3, green**). Induction of ATF3 was also not prevented or reduced by combined treatment with an anti-inflammatory and local anesthetic (**Figure 3, yellow**). These data suggest that inflammation is also not necessary for induction of ATF3 in sensory neurons innervating damaged skin.

**Figure 3.**
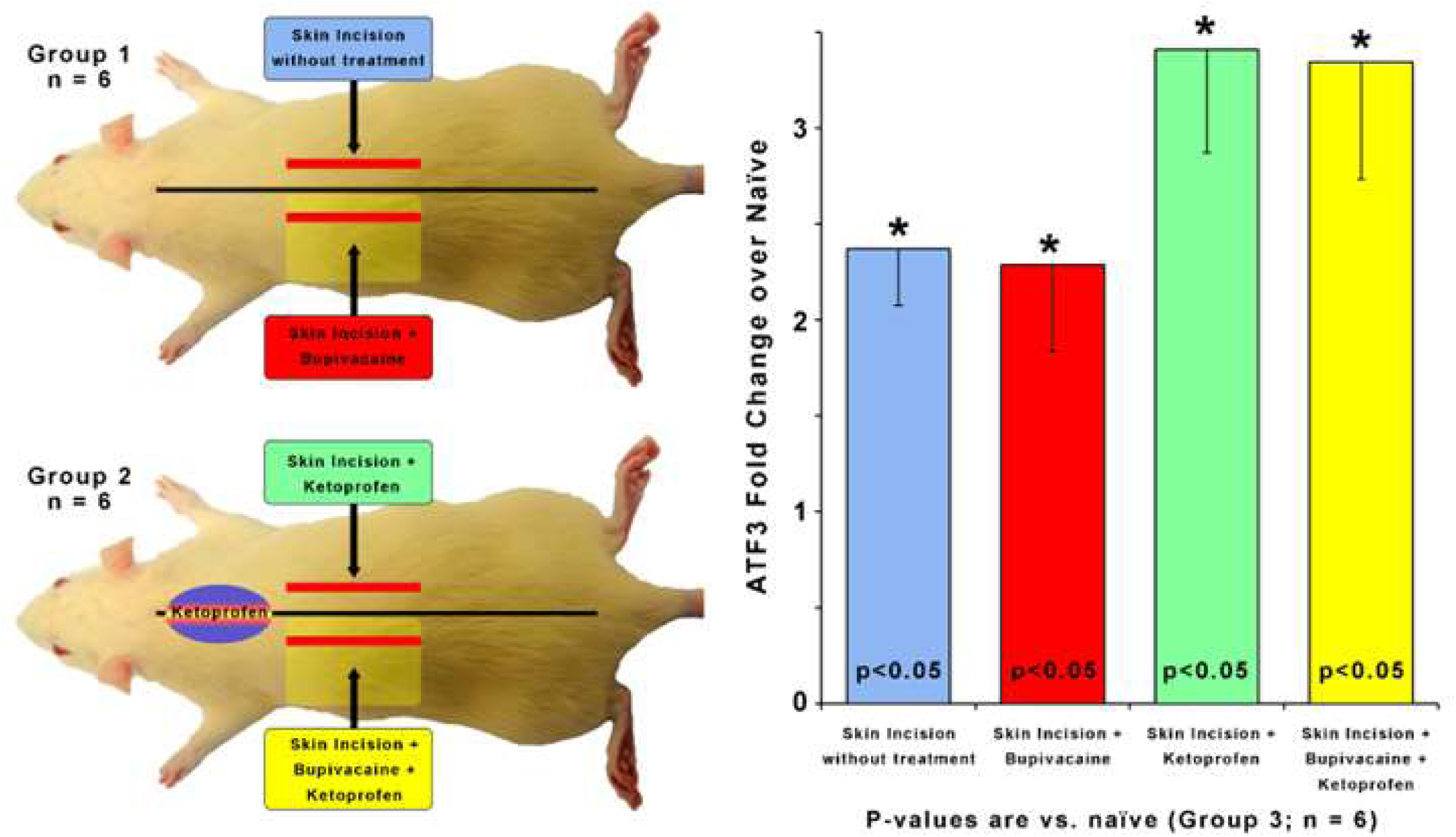
To determine if inflammation is required for ATF3 expression, a skin incision was made after injection of an anti-inflammatory (Ketoprofen; 10 mg/kg). Incision with/without local anesthetic (Bupivacaine; 0.125% in dH20) was also tested. ATF3 mRNA expression is shown on the right. P-values are for post-hoc t-test vs. naive. **Clinical relevance:** Ketoprofen is used peri-surgically to suppress the inflammatory response and reduce surgical pain. Despite the administration of Ketoprofen, or both Ketoprofen and Bupivacaine, ATF3 expression is still induced in DRG housing sensory neurons innervating the incised skin.

### Repeated tissue damage induces functional changes in a nociceptive reflex

We examined the behavioral response to repeated tissue damage compared to a single occurrence of tissue damage. We employed the well-characterized plantar incision model (PIM)[26, 27, 34, 36–47]. Mechanical nociception, assessed with von Frey filaments, revealed the expected reduction in threshold over the expected duration and the expected recovery of the threshold back to baseline (**Figure 4**). In the main experimental group (PIM/PIM), a second incision was performed as close to the same site as possible 4 weeks after the initial incision. The maximum threshold reduction in the 2-incision group matched that of the control groups that received a single incision at the same time and part of the same surgical batch (PIM/No Incision; No Incision/PIM). The temporal pattern of recovery after the first incision for the PIM/PIM group appeared not to differ from the PIM/No incision group. This indicates no overt difference in the background status of the PIM/PIM animals from the PIM/No Incision controls. Although the pattern of recovery of nociceptive withdrawal reflex threshold did not differ between the PIM/PIM group and the PIM/No-Incision group for the first incision, it differed significantly between the PIM/PIM group and the No Incision/PIM group after the 2^nd^ incision. The threshold of the PIM/No Incision group showed no changes from baseline during the same testing period, indicating there was no time-related change to the withdrawal reflex threshold after single incision, suggesting that the pattern of threshold change and recovery in the PIM/PIM group after the 2^nd^ incision was due, at least in part, to the history of prior incision on some element(s) of the reflex.

**Figure 4.**
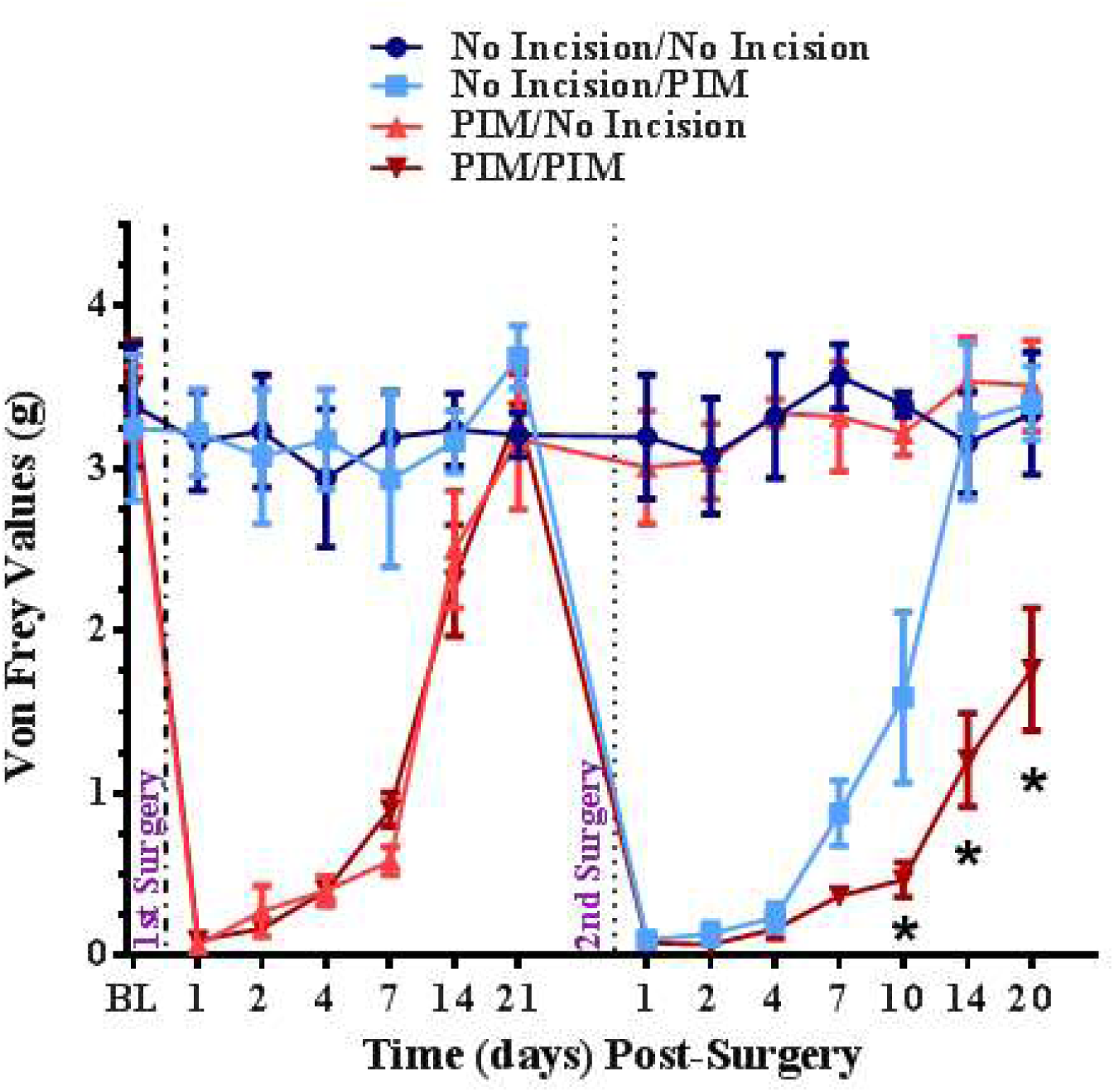
Changes in withdrawal reflex threshold over time after single or repeated incision. Surgeries are indicated by vertical dotted lines. *=p<0.05 RM-ANOVA and post-hoc t-test. **Clinical relevance:** Repeated injury is an etiological factor for persistent pain. Repeated injury in this clinically-relevant animal model induces a different response than single injury. Animal models of repeated tissue damage may be suitable for identifying mechanisms of persistent pain.

### Repeated tissue damage induces unique responses

Single incision results in a significant increase in ATF3 mRNA and protein expression in DRG innervating incised skin [10]. This increase abates somewhat over time from the initial maximum, but remains a highly significant increase even out to 28 days post-incision (**Figure 5 inset**).

**Figure 5.**
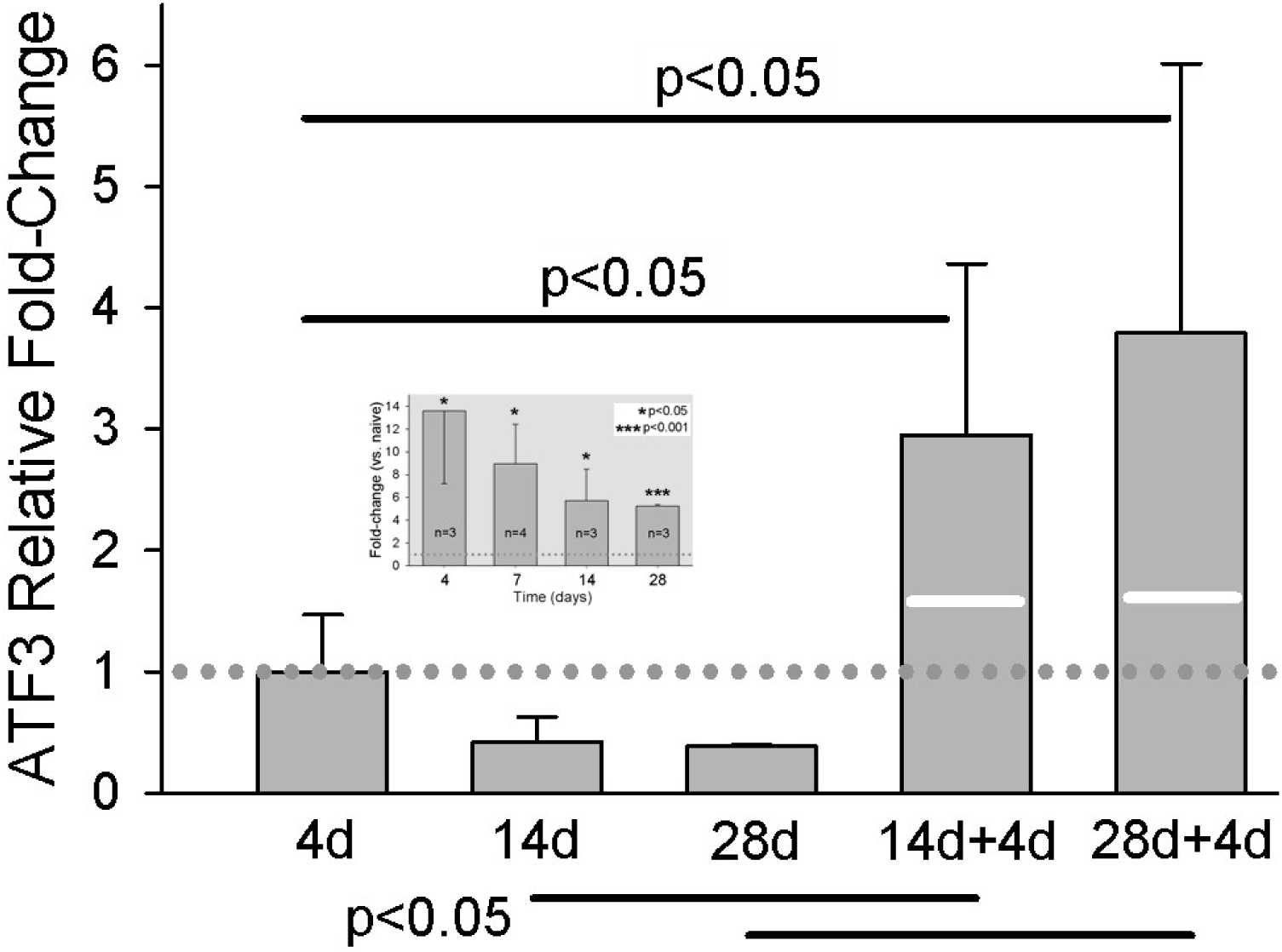
qPCR results for ATF3 at different times after single skin incision, or two different delays before a second skin incision. 14d+4d indicates: 1^st^ incision, 14 day delay, 2^nd^ incision, 4 day delay, euthanize. Similar for 28d+4d. Values of bars are fold-change vs. naïve, normalized against the mean of the fold-change at 4d (dotted line). White lines on the bars of the 2-incision group indicate the expression level that would be expected by simple addition of the levels from the same time points after single incisions. Data for single incision are the same as those presented in **inset**. N=7 for the 2-incision groups. Statistical analysis was ANOVA and post-hoc t-test. P-values in the figure are for the comparisons indicated by the bars.

Because one of the greatest predictors of persistent/chronic pain is repeated injury [48], we examined the effect of repeated incision on expression of ATF3 in DRG innervating repeatedly-injured skin. We hypothesized that ATF3-expression after skin incision might display conditioning characteristics as it does with repeated nerve injury [49, 50]. We induced a second incision, as close to the original incision as possible, at either 14 or 28 days after the original incision. We then took tissue 4 days after this second injury. The rationale was that this timepoint was the apparent peak of ATF3 expression after a single incision, so we chose to sample the presumable peak response of the second incision on the background of the first incision. Repeated incision induced an expression of ATF3 that was dramatically greater than for any single-incision group, and greater than the additive expression of the single-incision groups (**Figure 5**). These data suggest that ATF3 expression may indeed display a conditioning response to repeated tissue injury.

Qualitatively, it appears that the increased expression of ATF3 at the mRNA level is attributable to neuronal expression of ATF3 (**Figure 6**). Much like with single incision, we did not observe ATF3 immunohistochemical signal outside of neuron-like profiles. The increased ATF3 signal likely includes expression by more neurons, but certainly could include more ATF3 expression per neuron as well. We made no effort in these assessments to quantify either neuron number or degree of protein expression overall or per cell.

**Figure 6.**
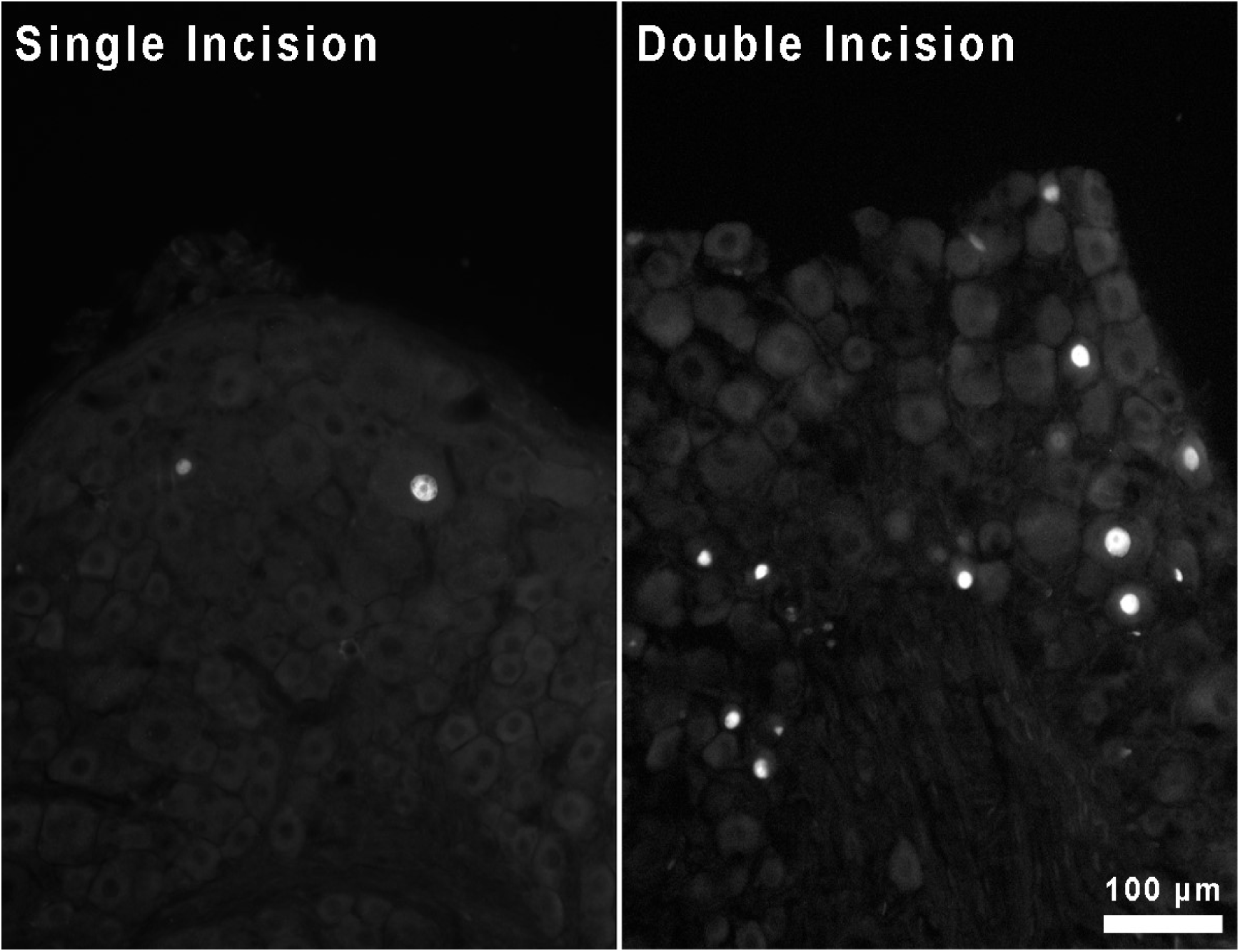
Immunohistochemical staining for ATF3 protein in sections of DRG housing neurons innervating skin incised either once or twice.

**Clinical relevance:** Repeated injury to the same region of tissue results in a unique temporal pattern of ATF3 expression in DRG housing sensory neurons innervating the injured tissue.

### Bioinformatic analysis identifies pain-related genes with ATF3 binding-sites

Nerve injury alters expression of ion channels, some of which are associated with pain. An extensive search of PubMed revealed no empirical evidence for ATF3 directly regulating expression of any ion channels in any setting. Although there is no direct evidence, it is clear that expression of both ATF3 and some ion channels is regulated in some of the same conditions. In order to focus our search for relevant genes, we sought to determine if any known pain-related genes had structures suggesting that ATF3 may play a role in regulating their expression. We examined the Molecular Signatures Database (MSigDB; Broad Institute; Human Motif gene sets – transcription factor targets) populated by the TRANSFAC data (v7.4). We examined the human genes having at least one occurrence of the following highly conserved motifs in the regions spanning 4 kb centered on their transcription starting sites [-2kb, +2kb]: CBCTGACGTCANCS = 257 genes, TGACGTCA = 235 genes, TGAYRTCA = 551 genes. The total number of genes with promoters containing at least one of these ATF3 consensus binding motifs was 671. We combined the results of three gene sets (1 for each of 3 separate ATF3 binding sequences) and cross-referenced this list against a set of Gene Ontology terms related to nociception or pain and against PubMed terms for pain. Of the genes with an ATF3-binding sequence, only 18 also had pain-related annotations (**Table 1**).

**Table 1.** Intersectional results of genes with both ATF3-binding sites and GO term annotations relevant to pain.

| ATF3-Binding Sites |  | Relevant GO annotations |  |  |
| --- | --- | --- | --- | --- |
| Genes | # sites | regulation of membrane potential | response to pain | sensory perception of pain |
| ADAM11 | 2 |  | YES |  |
| CALCA | 1 |  | YES | YES |
| BNIP3L | 3 | YES |  |  |
| CACNA1G | 2 | YES |  |  |
| CALM1 | 1 | YES |  |  |
| CAMK2D | 2 | YES |  |  |
| CFTR | 1 | YES |  |  |
| FGF12 | 1 | YES |  |  |
| GRIA4 | 1 | YES |  |  |
| GRIN1 | 1 | YES |  |  |
| HCN4 | 1 | YES |  |  |
| KCNA5 | 2 | YES |  |  |
| KCNF1 | 2 | YES |  |  |
| KCNN2 | 2 | YES |  |  |
| P2RX3 | 1 | YES |  |  |
| PHOX2B | 2 | YES |  |  |
| PLN | 1 | YES |  |  |
| SCN3B | 2 | YES |  |  |

### Pain-related genes with ATF3 binding-sites demonstrate unique expression after repeated incision

Skin incision is associated with significant electrophysiological changes in the ATF3-expressing sensory neurons [10], leading us to prioritize consideration of the 6 genes for voltage-gated ion channels. We chose to examine the expression of two genes known to influence electrical signaling in sensory neurons, particularly the depolarization phase of the action potential – Scn3b (Voltage-gated sodium channel beta subunit 3) (https://www.ncbi.nlm.nih.gov/gene/55800) and Cacna1g (Cav3.1 / T-type low-voltage-activated Calcium channel) (https://www.ncbi.nlm.nih.gov/gene/8913).

Scn3b mRNA expression appeared to be unaffected during the 4 weeks following the first incision (**Figure 7**, white bars). However, Scn3b expression was significantly increased after the second incision (**Figure 7**, black bars).

**Figure 7.**
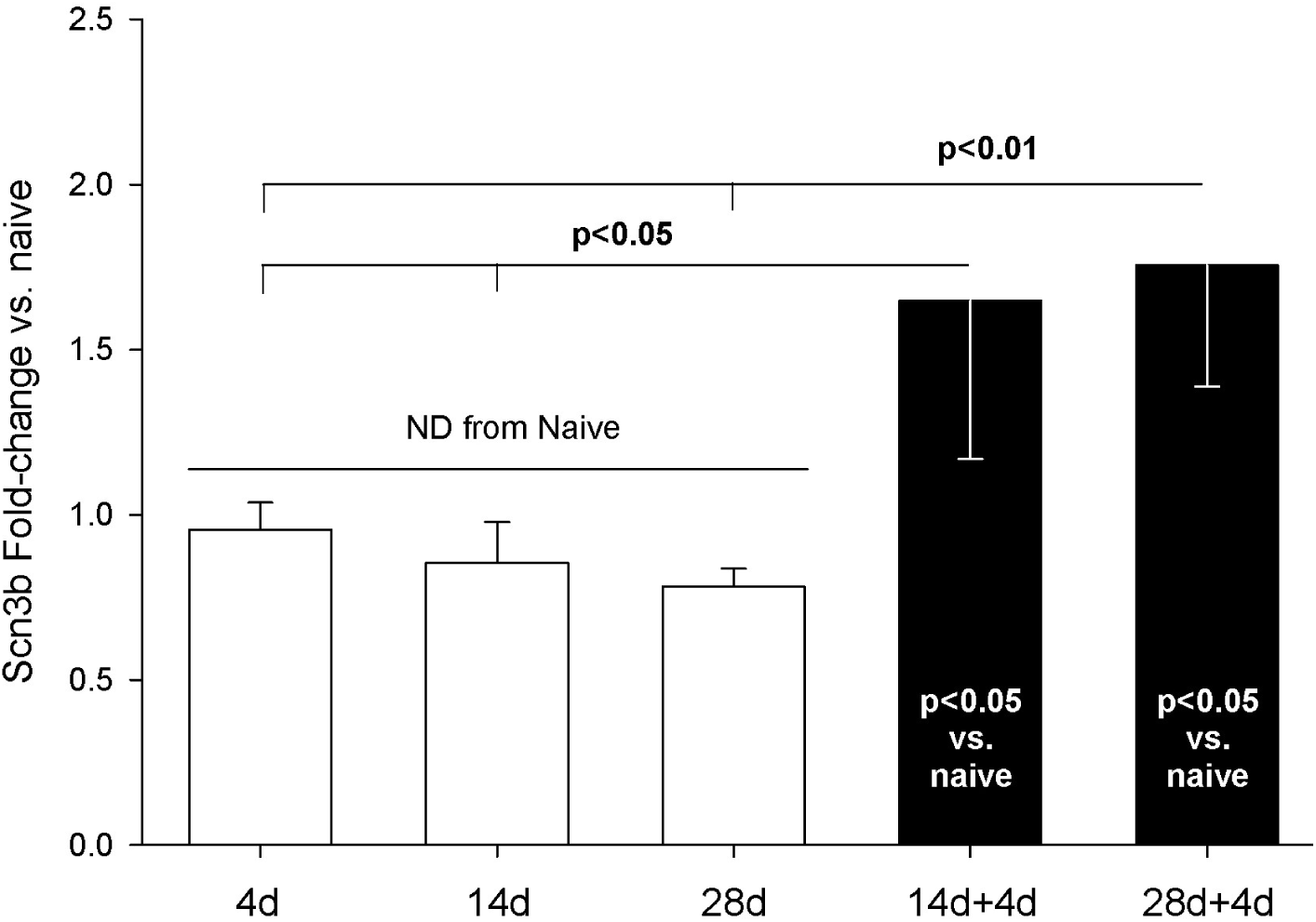
qPCR assessment of Scn3b mRNA from DRG housing sensory neurons innervating skin incised once (white bars) or twice (black bars). Single incisions were 4, 14, or 28 days earlier. Naïve (n=4), 4d (4), 14d (4), 28d (4), 14+4 (7), 28+4 (7). Statistical test was ANOVA and post-hoc t-test. P-values are from t-test. Scn3b expression was normalized to GAPDH. All groups were normalized to mean of Scn3b-v-Naïve, which was set to “1”.

Scn3b was identified by screening genes with ATF3-binding sites for pain-related annotations. We therefore examined whether there may be a relationship between expression of ATF3 and Scn3b on an animal-by-animal basis across all groups. There was a significant positive relationship between expression of ATF3 and Scn3b (each vs. GAPDH) (**Figure 8**).

**Figure 8.**
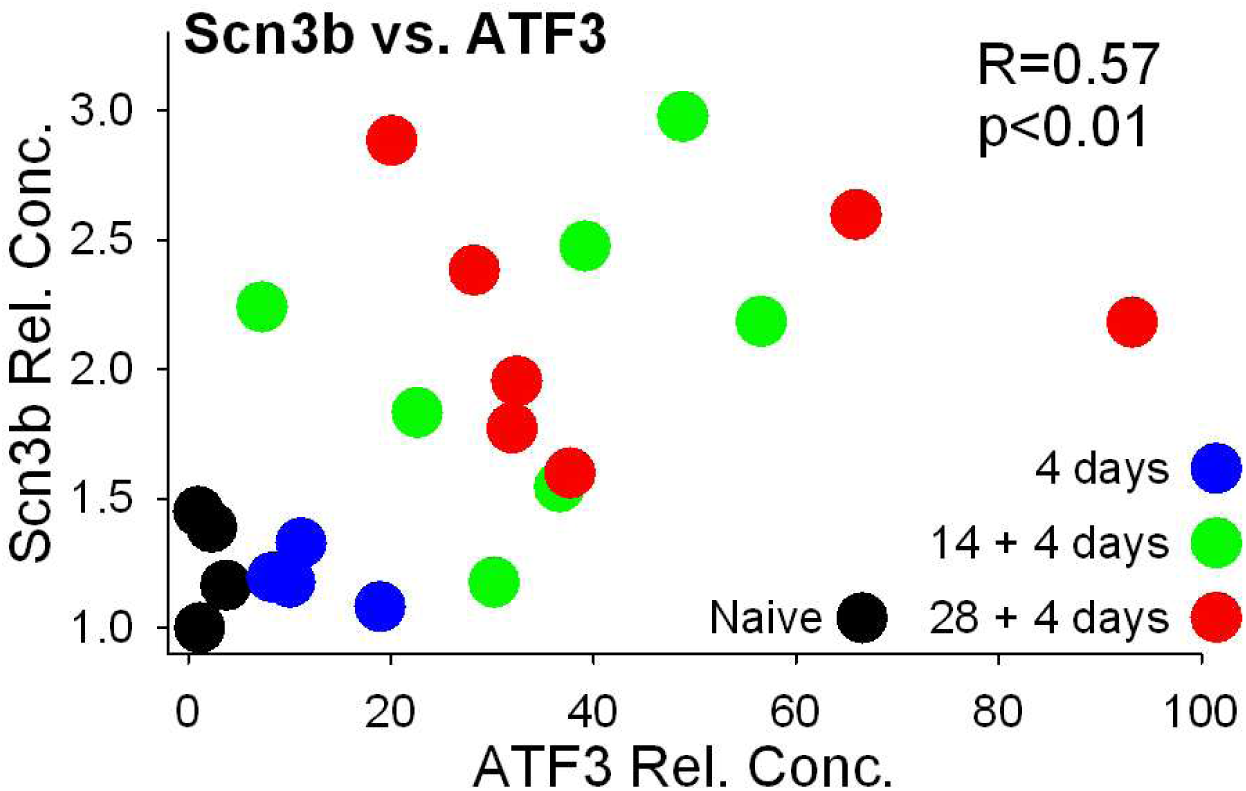
Expression values of Scn3b and ATF3 (both normalized to GAPDH). Values were compared by Pearson correlation.

Unlike Scn3b mRNA, expression of Cacna1g mRNA was significantly reduced by 4 days after a single skin incision (**Figure 9**). This significantly-reduced expression was reversed 4 days after a second incision. Expression after repeated incision was no different from naïve, while after single incision it was significantly reduced.

**Figure 9.**
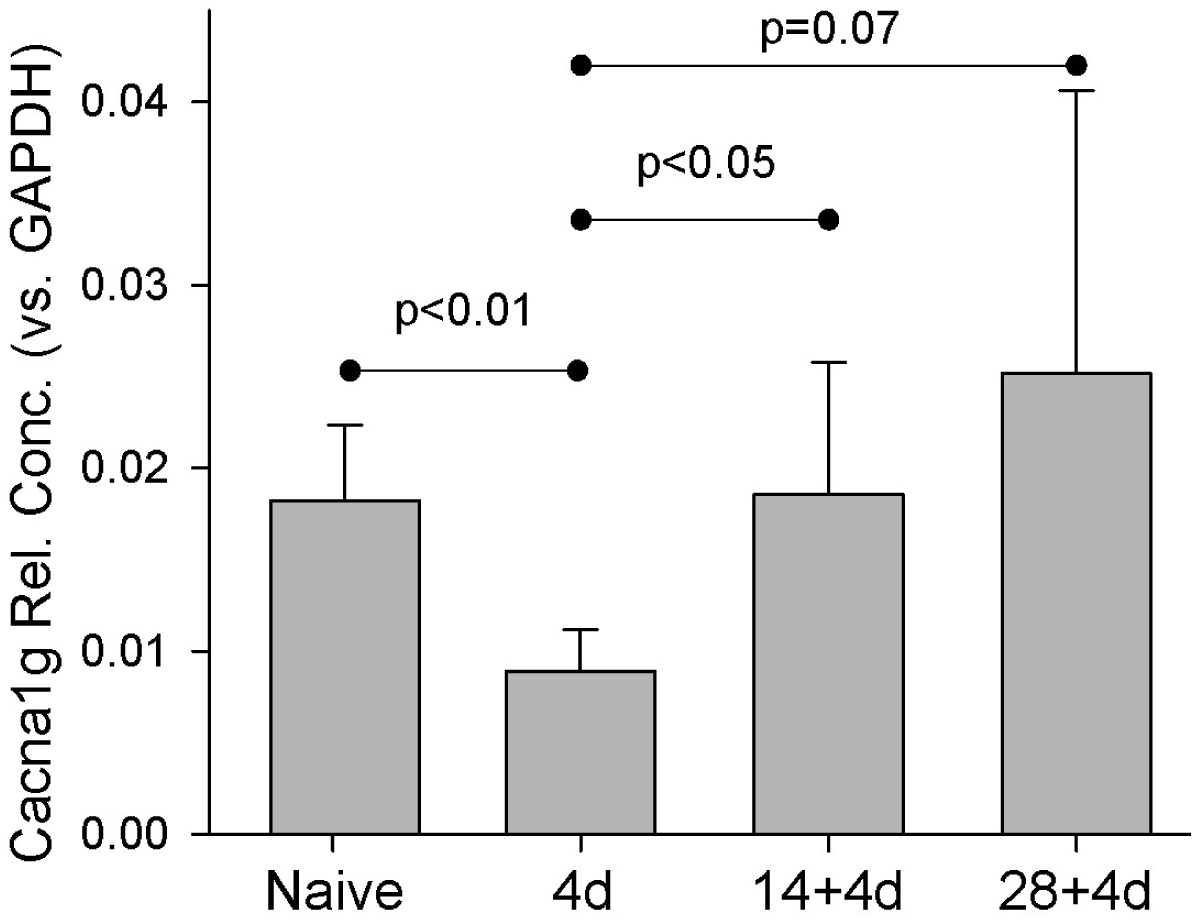
Naïve (n=4), 4d (4), 14d (4), 28d (4), 14+4 (7), 28+4 (7). Statistical test was ANOVA and post-hoc t-test. P-values are from t-test. Cacna1g expression was normalized to GAPDH.

We then compared the levels of each of the genes within each animal. Because expression of each gene was different between conditions of single- and repeated-injury, but was similar across the 2 repeated-injury groups, we combined animals of the 14+4 and 28+4 repeated-incision groups under the same label of “Double Incision”. These are compared those to animals of single incision and naïve groups (**Figure 10**). It is clear that the expression from naïve and single incision animals cluster together, while the expression from the double incision animals is much more distributed. Similar to the pattern of expression with Scn3b, expression of all 3 genes in the repeated-incision group is greatly separated from the single incision animals.

**Figure 10.**
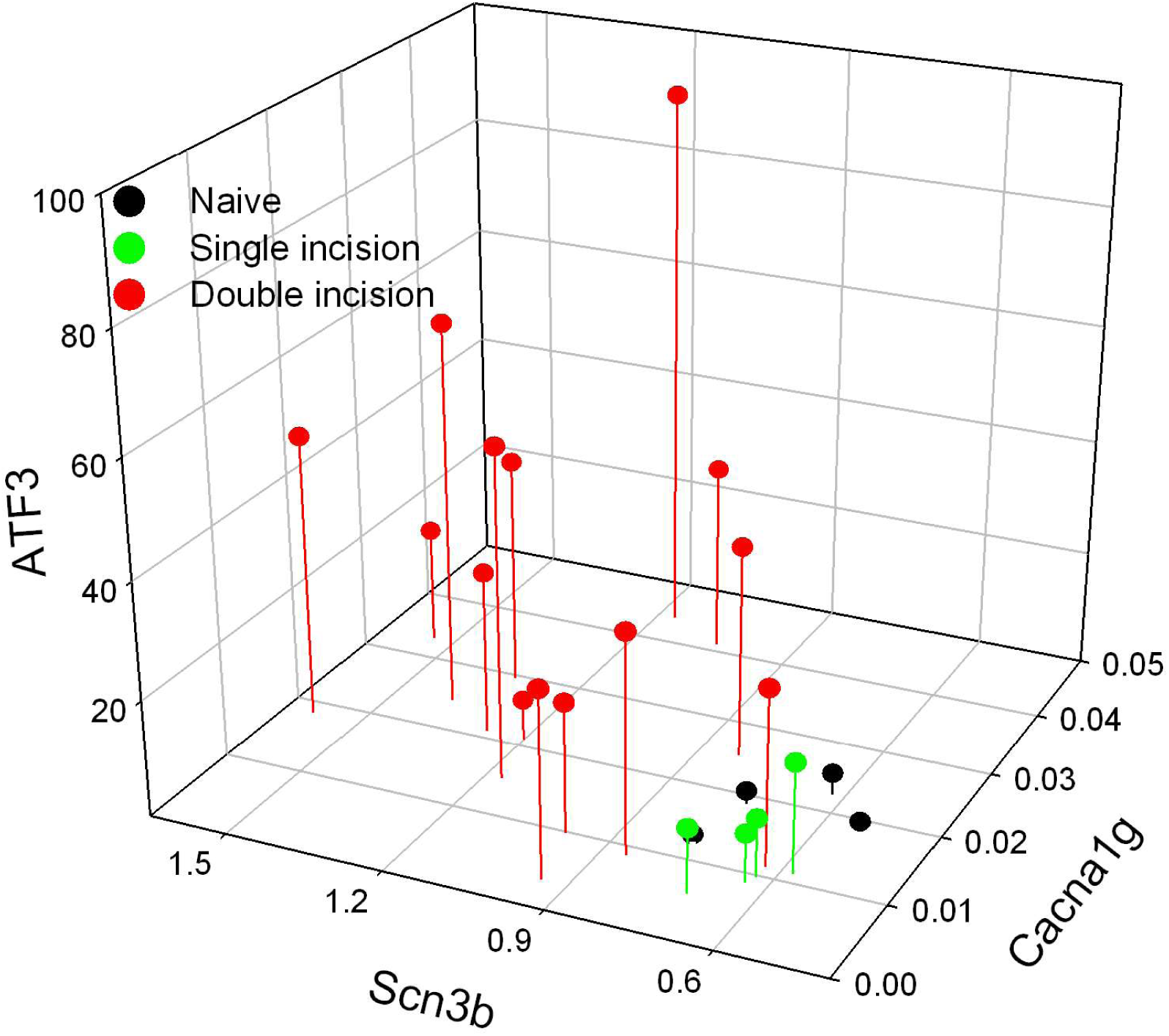
The relative concentration (vs. GAPDH) of each of the genes is plotted on a per-animal basis. Both groups of animals with two incisions (14+4 and 28+4) are labelled as Double incision. **Clinical relevance:** Effects on expression of pain-related genes differ between conditions of single and repeated tissue damage. These animal models suggest mechanisms by which repeated injury contributes to the etiology of, and risk for developing, persistent neuropathic pain.

### Wound closure method and treatment affect induction of ATF3 and GAP-43 after incision

Wound closure with suture or staples clearly results in changes in expression of genes in a manner similar to what occurs in nerve injury [9, 10]. Glue-closure is also used with some wounds [51–57]. We therefore examined the effect on gene expression of incision followed by wound-closure with surgical glue.

Similar to ATF3, growth-associated protein 43 (GAP-43, B50) is upregulated in sensory neurons in response to nerve injury (e.g., [50, 58, 59]) and after skin incision [9, 10]. As expected, ATF3 and GAP-43 expression was significantly increased 4 days after the incision was closed with surgical staples. Because many cutaneous wounds are closed with cyanoacrylate adhesive, we examined whether this closure method might result in a different outcome for the markers of axonal injury. These increases were significantly reduced when the wound was closed with surgical glue (**Figure 11**).

**Figure 11.**
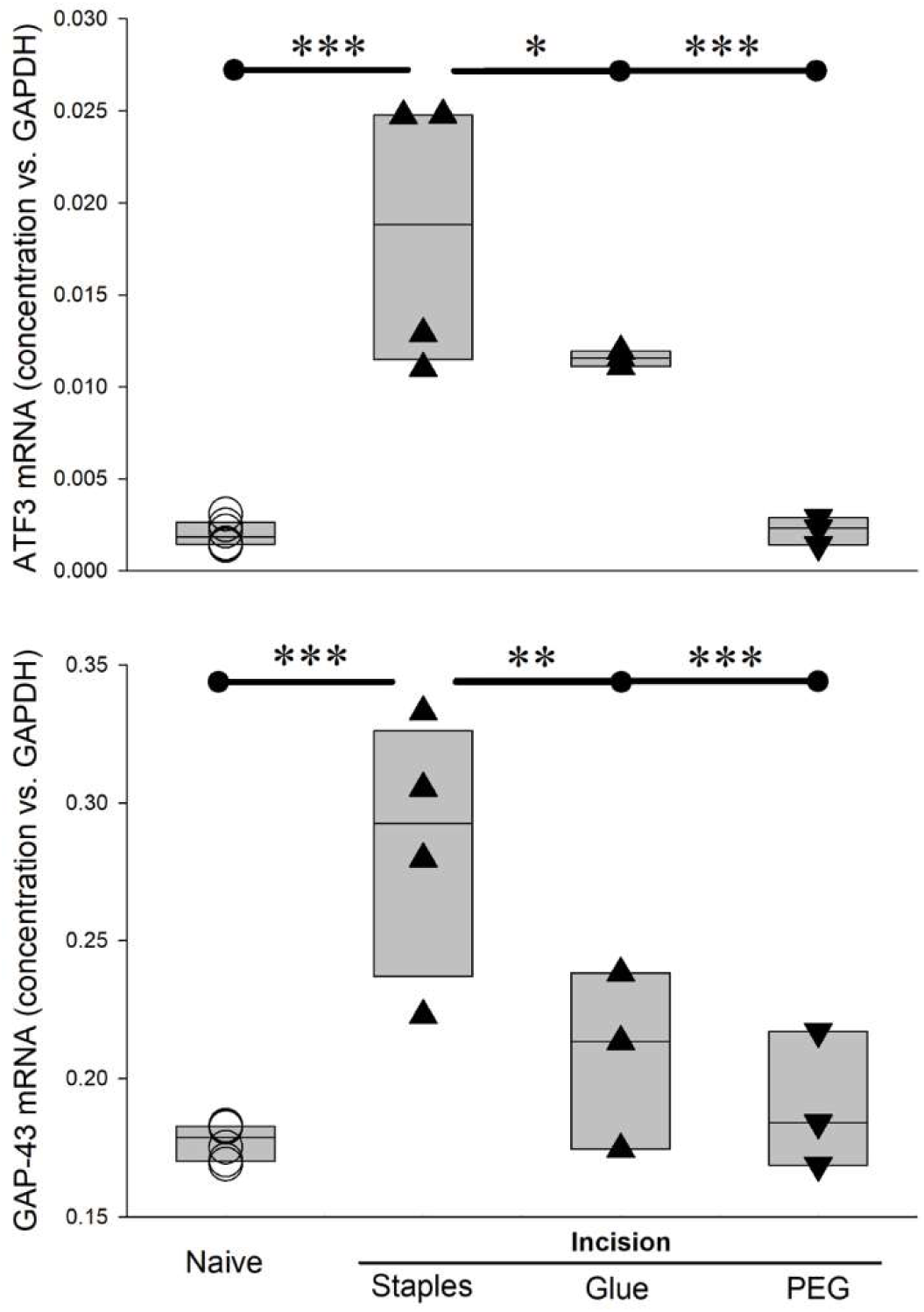
Different wound-closure methods, and post-incision treatment, affect ATF3 and GAP-43 expression differently 4 days after incision. *=p<0.05, **=p<0.01, ***=P<0.001 by ANOVA and post-hoc t-test with control set to incision+staples. Naïve (n=6), Incision+staples (n=4), incision+glue (n=3), incision+PEG (n=3)

Our working hypothesis has been that tissue damage, including surgical incision, induces damage to axonal integrity to a sufficient degree that the positive and/or negative centripetal signals indicating injury become sufficient to induce gene expression changes similar to those induced by overt damage of nerve tissue. Under this hypothesis, we examined whether a treatment known to maintain or restore membrane integrity, including axonal membrane, might influence these changes in gene expression. Polyethylene glycol and other “fusogens” have been used for decades by labs working from this idea of membrane repair or stabilization [21, 24, 25, 60–68]. The understanding of mechanisms underlying the effects of the fusogens has advanced significantly, even to the stage of enabling veterinary and human clinical trials [18, 19, 69–71].

We administered PEG acutely post-incision, applied topically to the incision site and subcutaneously as was done previously [25, 63] and closed the incision with staples. In rats treated with PEG, there was no upregulation in expression of ATF3 or GAP-43 mRNA at 4 days after incision (**Figure 11**), a time chosen because it appears to be the peak of the ATF3 response.

Following the promising early-stage results from PEG-treatment on expression of ATF3 and GAP-43 mRNA, we performed additional experiments to examine expression of ATF3 protein and functional properties of sensory neurons traced from the incision site, similar to our prior work [9, 10, 72–76]. We assessed some of the excitability-related electrophysiological properties of sensory neurons DiI-traced from the incision site or an equivalent site in naïve non-incised animals using patch clamp of single-neurons dissociated 28 days after incision. Following recordings, the cells were fixed and assessed with immunohistochemistry against ATF3. This provided a means to directly associate the ATF3 expression with electrophysiological properties at the single neuron level (**Figure 12A**).

Rheobase (depolarizing current threshold for inducing action potential firing; **Figure 12B, green trace; 12C bar graphs**) was significantly lower in ATF3+ vs. ATF3-in the incision group, and was the lowest (most-excitable) vs. all other groups, in agreement with our prior work [10]. Also as expected, the proportion of traced neurons that were ATF3+ was greater in the incision group (**Figure 13C, red/yellow pie charts**). Interestingly, the proportion of ATF3+ neurons was reduced in the PEG-treated group (**Figure 13D, green pie charts**), similar to the proportion from the traced/non-incised group (**Figure 13D, blue pie charts**).

As we reported previously [10], many of the ATF3+ neurons displayed repetitive firing in response to even low-level depolarizing currents (**Figure 13B, red trace**), which does not usually occur with acutely-dissociated sensory neurons. We quantified this response profile and determined that the ATF3+ neurons displayed significantly greater repetitive firing than ATF3-neurons (which had essentially no repetitive firing), regardless of group (**Figure 13D**). This significant difference occurred at much lower levels of depolarizing stimulation for the incision group than for the tracer-only groups and the incision+PEG group.

**Figure 12.**
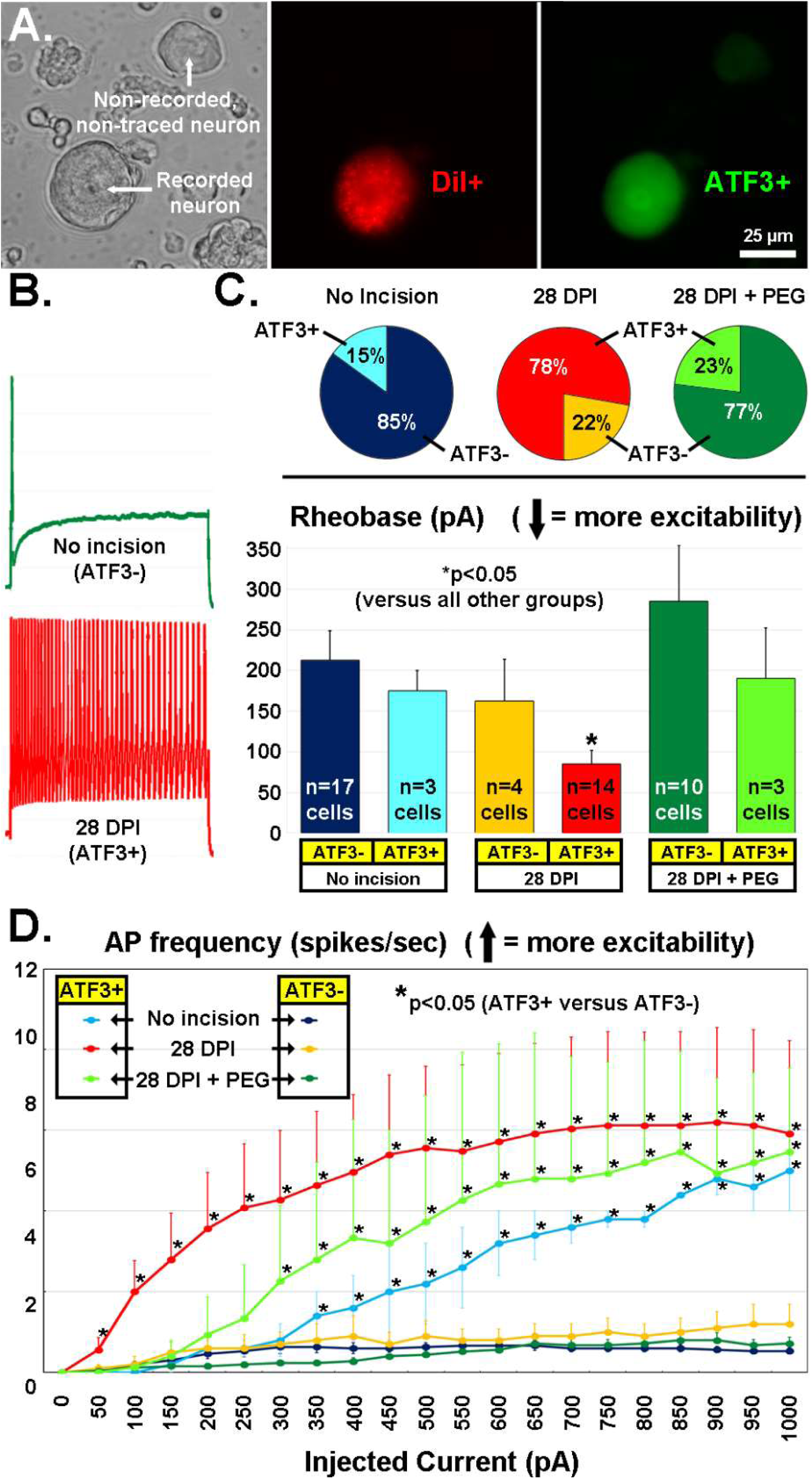
A) Recordings were made of acutely-dissociated DRG neurons (left panel, DIC image) labeled from the incision site with the retrograde tracer DiI (middle panel, red) and later run for ATF3 immunohistochemistry (right panel, green). B) Examples of firing pattern (in response to a 1s, 1000pA depolarization) from an ATF3-neuron from a naïve animal (green) and from an ATF3+ neuron traced from the wound site 28d after incision. C) Proportion of DiI-traced and recorded neurons that expressed ATF3 in the 3 different treatment groups (pie-charts). Rheobase current of DiI-traced and recorded neurons parsed by expression of ATF3 in the 3 different treatment groups (bar graphs). D) Number of APs fired (y-axis) during a 1s depolarizing pulse (x-axis). Error bars are SD. *=p<0.05 in (C) (vs. all other groups) and (D) (ATF3+ vs. ATF3-) **Clinical relevance:** Acute-stage application of PEG may prevent long-term injury-related changes in gene expression and electrophysiological properties, which might in-turn reduce persistent pain after tissue damage. There is also reason to suspect that glue-closure may provide better pain-outcomes than suture or staples for suitable conditions.

## DISCUSSION

It is clear that many people experience pain after tissue damage, including surgery. In many cases this pain persists beyond apparent wound healing and has signs of being neuropathic, including being resistant to anti-inflammatory treatments. Conventional perspectives on this kind of pain are failing, and both research and clinical practice are calling for novel perspectives and development/inclusion of more mechanism-based diagnosis/treatment and therapy-development (e.g., [6, 77, 78]). Here we extended prior work by our lab and others that consider the effects on sensory neurons of damage to peripheral target tissues as similar to the effects of nerve injury, a known major etiological factor for developing persistent neuropathic pain. We examined the influence of common clinical analgesic approaches on the nerve injury-like biological responses induced by tissue damage. We further examine the effects of repeated injury, which has great potential as an etiological factor for persistent neuropathic pain, but one that has not received much attention. Finally, we examined the efficacy of some feasible approaches to prevent the nerve injury-like response in sensory neurons.

We used the de novo expression of ATF3 mRNA as a surrogate for the larger functional effects that we described previously [10] in an effort to determine the necessary and sufficient factors for the response. Pre-surgical local/regional anesthesia and peri-operative anti-inflammatory regimens are common approaches for reducing peri-operative pain, but they have unclear value for reducing the emergence of persistent pain [6, 79]. For the injury/stress response induced in sensory neurons to be considered a feasible contributor to persistent pain as we propose, it should also be resistant to any treatments that do not clearly prevent emergence of persistent pain. ATF3 was expressed despite a clinically-modelled regimen of the anti-inflammatory ketoprofen, indicating both that post-incision inflammation was not required for the response and suggesting that standard clinical anti-inflammatory approaches are not likely to affect this response (**Figure 3**). ATF3 was also expressed despite a clinically-modelled application of local anesthetic, indicating both that AP conduction from incision site to DRG was not required for the response and suggesting that standard clinical local/regional anesthetic approaches are not likely to affect this response (**Figure 1**). These findings alone provide a strong validation for this response as a potential contributor to clinical persistent post-surgical pain as the response appears resistant to the most-used approaches to prevent or reduce post-surgical pain.

The magnitude of ATF3 expression after skin incision might be considered relatively small. On the surface, this might suggest that the effect of so few neurons expressing ATF3 may be minimal, especially comparing the amplitude in response to that with nerve injuries. However, as discussed in our prior work [9, 10] and supported by the findings here of the requirement for axonal damage (**Figures 2, 13**), the relatively small magnitude is much more a function of the small proportion of neurons in each DRG that is involved in the tissue damage. However, even a focal and small injury can have effects that become fairly widely distributed anatomically and functionally. For example, other nerve injury-like responses may occur in response to single or repeated tissue injury. If so, there may also be responses such as macrophage infiltration and migration to the ATF3+ sensory neurons as well as microglial activation around the dorsal root projections in the spinal cord. Both of these contribute to pain mechanisms which can sensitize neighboring neurons that were not themselves injured, again expanding the effects. It is also possible – however inelegant – that there may be a “bulk effect” determining the relationship between ATF3 expression and the emergence of pain. The number or proportion of sensory neurons in which ATF3 expression has been induced as a result of some event or condition may indeed be a controlling factor in the emergence of neuropathic pain [80–82].

Although we did not set out to test the threshold for what noxious stimuli are required to induce ATF3 expression in sensory neurons, the study nonetheless offers some insight. The dermatome-mapping process itself – inducing the CTM reflex by noxious pinch – was not sufficient to induce detectable ATF3 expression. (**Figures 1-3**). For example, the mapping process for **Figure 2** definitely triggered T11 axons, but there was no ATF3 detectible in the T11 DRG. This is consistent with another study we performed which applied many noxious pinch stimuli yet observed no ATF3 expression [83].

**Figure 13.**
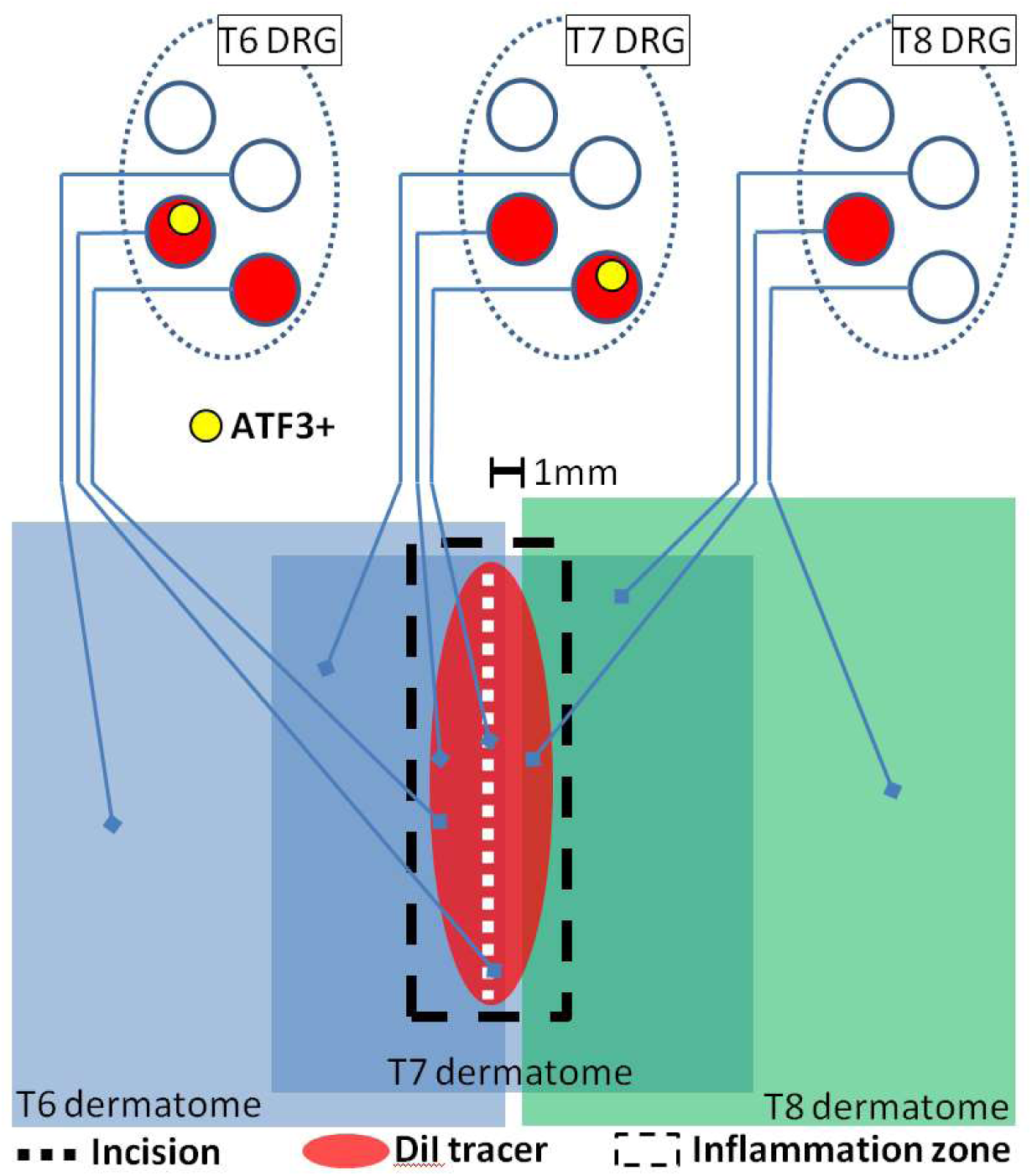
Schematic representation of presumed interaction of cutaneous neuroanatomy, retrograde tracer, and incision-associated effects. The figure incorporates conclusions from the experiments reported in Figures 2, 3, and 13. Note that dermatomes are C-fiber dermatomes which overlap significantly, not the classical A-fiber dermatomes which do not overlap.

### Considering axonal injury in skin versus nerve

Much of our rationale for proposing that this response in sensory neurons to tissue damage should be considered as a possible contributor to persistent post-surgical pain is due to its similarity to the sensory neuron response to nerve injury, which is a proven etiological factor for persistent neuropathic pain. In this context we should consider how the core injury – that of injury to the axon – might differ in the setting of skin vs. nerve. The full neural “cell body response” to nerve injury requires both positive/injury signals (such as certain cytokines) and negative/absence signals (e.g., lack of constitutive retrogradely-transported target-derived factors) [84–88]. Injury to a discrete peripheral nerve induces positive/injury signals directly in the injured nerve tissue and results in negative/absence signals by virtue of disconnecting the biochemical transport pathway from target tissue to soma. Injury to target tissues induces positive injury signals that are presumably retrogradely transported similar to what occurs with nerve injury, but it is possible that because the axons are still resident in their target tissue they might not convey an absence signal (i.e., the axons may still have access to those target-derived signals and retrogradely transport them). That is, target-derived positive/intact signaling – both electrical and/or biochemical – may remain at a level that is sufficient to either entirely or partially prevent the positive injury signal from inducing a cell body response. It is this possibility that was tested by the experiments made at progressive distances outside of a defined receptive field/dermatome (**Figure 2**). The conservative perspective was to hypothesize that the positive injury/inflammation signals induced by the incision 1mm outside the dermatome would be capable of inducing ATF3, and the incision at 5mm outside would not. But this did not occur, suggesting that the level/type of positive injury/inflammation signal induced by incision used here was not capable of inducing ATF3, implying that overt injury to the tissue-resident axons is required for induction of ATF3 expression.

This is not to imply that inflammation alone would not be capable of inducing a nerve injury-like response. Clearly, inflammation of peripheral target tissues can induce many of the same effects we have described in terms of sensitization of electrophysiological properties/responses and nociceptive behavioral responses. However, those are largely prevented by anti-inflammatory treatments and it appears that only the most severe manifestations of inflammation – potentially non-physiologic – and chemical stimuli which may themselves result in damage to the tissue or the responsive axons are capable of inducing ATF3 expression in the sensory neurons innervating the inflamed tissue [89].

The distinction between axonal injury associated with target tissue damage and nerve damage may appear esoteric, but it is not. First, the Gross Anatomy and Histology perspectives can provide real clinical and mechanistic meaning. In terms of sensory neurons, it has been recognized for over 100 years that injury to dorsal root and to peripheral nerve result in different sensory neuron responses, with more recent studies providing some molecular and cellular signatures [49, 90–92]. It may be that there is yet another different response when axons are injured within the setting of their target tissue and the wide range of distinct cellular compositions. Single axons of sensory neurons traverse many different tissues, each of which has a distinct cellular composition. Dorsal root, DRG, nerve, and target tissue have different resident immune and supporting cells, and this composition may become altered differently by injury to each of these tissues.

Second, Schwann cells are intimately associated with PNS axons across all of the tissues through which nerves traverse. It is increasingly recognized that there is a range of Schwann cell types which differ in accord with the axonal composition of a nerve (sensory or motor) and their central/peripheral and tissue location [93–95]. Terminal Schwann cells differ with the tissue/structure innervated [96, 97]. Axonal injury induces differentiation of a “Repair Schwann cell” phenotype [98–101]. The interaction between damaged axons and their associated Schwann cells may therefore differ by location of injury.

Third, outside of an overt injury to a distinct peripheral nerve, there is little recognition of neuropathic mechanisms when considering potential factors contributing to tissue damage- or surgery-associated persistent pain. If neuropathic mechanisms are considered outside of overt nerve injury, these are often characterized as idiopathic or covert injuries to nerves, largely as a diagnosis of exclusion when anti-inflammatory treatments fail. In these cases, the field looks to any number of CNS mechanisms. However, it is possible that tissue damage from any range of sources results in concurrent damage not to discrete peripheral nerves, but to tissue-intrinsic axons, and that the axonal damage triggers a cellular injury response in sensory neurons that resembles that induced by nerve injury. Such a neuropathic mechanism would require therapeutic approaches appropriate for the PNS, which often differ from those required for CNS pathologies.

### Considering repeated damage

It is uncommon that a single incident of tissue damage, surgical or otherwise, results in persistent neuropathic pain. Certainly there are reports that suggest this in the literature for conditions such as CRPS, though it is becoming increasingly recognized that labelling the precipitating event as a “single non-notable injury” may be misleading and incomplete, as there may be predisposing factors that are not readily apparent in the medical history [102–104]. It is a fundamental principle of biology that prior experience can influence the response to later experience. Relevant here is that there is suggestion from both clinical experience and basic science that repeated injury can change and/or amplify the response, including from the most common surgery performed globally – Cesarean section – from which chronic post-surgical pain is not rare (e.g., [49, 88, 105–116]).

We therefore examined whether the response to repeated injury might differ from single injury. The response to repeated injury assessed behaviorally indicated a longer-lasting hyperalgesia/sensitization response than for single incision. Amplitude could not be examined here because the extreme sensitivity of the withdrawal reflex after even a single incision created a floor-effect. It is also unclear how long the reflex might remain sensitized after repeated injury.

The effect of repeated injury assessed at the transcriptional level indicated unique responses. Expression of ATF3 was increased super-additively or synergistically in response to a second incision. It is not clear if this greater signal in bulk mRNA was due to the same cells expressing more ATF3, the same level of ATF3 expressed in more cells, or both. This detail will need to be clarified as the transcription-regulation actions of ATF3 are context-dependent based on cell type, condition, available binding partners, and the level and time course of expression (e.g., [117, 118]). At the cellular/tissue level, it is not clear if the unique effects of repeated injury is due to an altered inflammation in repeatedly-injured tissue, or perhaps because the initial injury induces a higher innervation density so that a second injury might affect more sensory neurons.

#### Scn3b

Although the details are not entirely clear [119], Scn3b does have the capacity to influence the current density, activation threshold, and inactivation kinetics of voltage-gated sodium channels, including those with alpha subunits Nav1.3, Nav 1.7, and Nav1.8 (e.g., [120]). Scn3b is upregulated in Chronic Constriction Injury, nerve injury, and streptozocin models [121–123], all of which also induce expression of ATF3 in sensory neurons [124–128]. Scnb3 is upregulated in our model uniquely after repeated injury. It is not clear if this substantially-increased expression is in the same cells expressing Scn3b previously (small diameter sensory neurons [121]) and/or additional cells, such as the large-diameter sensory neurons that normally express only Scnb1.

#### Cacna1g

“T-type calcium channels are uniquely… “first responders” to depolarization. The low voltage threshold for activation of T-type channels drives their opening in response to relatively small positive changes in membrane potential” [129]. T-type channels are involved in low-threshold calcium spikes, neuronal oscillations and resonance, and rebound burst firing, all of which have been associated with a range of painful conditions. Even small fluctuations in voltage mediated by these channels means that there may also be a biochemical signal because the current carries calcium, a powerful signalling molecule – “…[T-type calcium channels] contribute to regulating intracellular calcium levels near the resting potential of many cells” [129].

Cacna1g/Cav3.1 mRNA is downregulated by single incision, but returns to pre-incision levels with repeated incision. The exact nature of the unique response of Cacna1g/Cav3.1 is not entirely clear, because we do not yet know how it is regulated at 14d and 28d after single incision. It is possible that expression continued to decline or returned to pre-incision levels. Regardless, the 14+4 and 28+4 responses differ meaningfully – the prior incision changes the response to the second incision.

We performed assessments of nocifensive behavior out to 28d after a second incision. However, the molecular assessments were made only out to 4d after a second incision. Although these data are strongly suggestive of a unique transcriptional response to repeated incision, a more detailed examination of the molecular expression pattern, especially in conjunction with behavioral pharmacology and electrophysiological assessments, would provide a better indication of the mechanisms at play.

We did not perform an analysis of the withdrawal reflex beyond 4 weeks after the 2^nd^ incision. The trajectory of threshold recovery suggests that the recovery is likely to be extended in duration. It is also possible that it may not return to baseline, unlike what occurs with single incision. The Taylor lab has demonstrated that there is a latent sensitization even after single insults, the masking of which relies on opioid and NPY signaling [38, 40, 130–132]. Repeated injury may also influence the masking mechanisms. It is very enticing to consider the possibility that the masking and sensory axon injury-induced sensitization mechanisms may intersect and provide a clinically-relevant mechanism for persistent pain.

### Co-regulation with ATF3

Our molecular assessments were targeted toward genes that might be regulated by ATF3 and might have a role in the long-term electrophysiological sensitization we have observed. We therefore considered whether there might be signs of co-regulation. It is very clear that ATF3 and Scn3b were strongly similarly-regulated, and the expression levels for DRG from repeated injury grouped well outside of the expression levels from single injury. Further, despite the fairly variable expression of Cacna1g, it contributed strongly to a picture of consistent co-regulation of the ion channel genes with ATF3 at the animal level. Determining whether ATF3 directly regulates expression of Scn3b, Cacna1g, and the other genes in Table 1 could be very informative for understanding how tissue damage leads to persistent pain-related changes in sensory neurons but would require more refined assessments such as performing similar assessments using ATF3-null mice.

These data support the proposition that tissue damage induces neuropathic-like responses in sensory neurons in addition to the well-established responses to inflammation condition, either or both of which may contribute to persistent pain conditions. The expression of ATF3 in sensory neurons may serve as an experimental biomarker which can be used to assess a range of chronic pain models for possible contributions due to sensory neuron injury and neuropathic response. Despite ATF3 being induced by both nerve injury and tissue-damage/skin-incision, it is not yet clear to what degree the overall genetic responses are similar or different. Our data indicate that the GAP-43 response to skin incision is also similar to nerve injury, but the response of Cacna1g and Scn3b are not as clearly similar. It is possible that tissue injury will only partially reproduce the effects of nerve injury, and/or that repeated tissue damage may be more similar to nerve injury than a single incidence of tissue damage. These possibilities can be readily addressed with transcriptomic approaches.

Because there currently is little recognition that “simple” tissue damage, especially when it “heals normally”, might be relevant to medical history for pain, it will be very difficult to retrospectively construct an accurate estimate of the potential contribution of this response, due to single or repeated injury, to persistent pain. Instead, it must be determined prospectively, making it important for the field to consider this biological response alongside other potential contributing factors moving forward.

### Considering the response among other known processes and factors

The biological response described here could provide mechanisms contributing to currently unknown and/or poorly understood etiological factors for persistent pain after tissue damage. It is clear that not all incisions or tissue damage incidents are chronically-painful or sensitized, despite the responses we present here appearing to be quite powerful and consistent. Clearly these responses alone do not dictate outcomes but are instead working in conjunction with other responses and systems.

If the “bulk effect” factor does play a role in the emergence of persistent pain, conditions with repeated injury may become more important. Neighboring non-injured axons will respond to the tissue damage-induced inflammation (which includes significantly increased production of NGF [34, 133–135]) by undergoing collateral sprouting [136, 137]. This could increase the number of neurons whose axons would be damaged by the second injury, ultimately increasing the number of neurons affected by the injuries.

It is currently unclear, but it is possible that the injury response in sensory neurons induced by tissue damage might induce similar responses in spinal cord and DRG as are induced by nerve injury such as activation of spinal microglia associated with central axons of the injured sensory neurons (e.g., [138]) and infiltration of macrophages into the DRG and localization near the soma of injured neurons where they can influence neighboring non-injured neurons (e.g., [139]). We also cannot yet be certain that the response we describe underlies the latent sensitization described by Taylor (e.g., [38, 40, 140]), but it seems both feasible and likely that it at least contributes. We have yet to determine how, and if, these responses interact with the range of known analgesic adaptive mechanisms. But it is conceivable that persistent pain could emerge if the adaptive mechanisms were to become saturated (e.g., if too many sensory neurons are involved) or if some individual variance (genetic, disease, etc.) made one or more of the adaptive mechanisms less functional.

An additional consideration is that splice isoforms have been described for Cacna1g, Scn3b, and ATF3 [141–143]. This means that the qPCR signal we report here might be reflective of expression of different isoforms instead of, or in addition to, regulation of the number of copies. Alternative splicing can introduce even greater functional plasticity than up- or down-regulation of a single type of transcript [144]. Cacna1g meets all of the rationally-constructed criteria for having functionally significant splice variants [145], and isoforms of ATF3 have been experimentally demonstrated to be expressed and exert different effects on transcription [142].

### Considerations for Glue-Closure

Not all wounds can be effectively and safely closed with adhesive, but glue-closure is used in many clinical settings and cyanoacrylate-type products are an integral part of the surgical arsenal. There have been many studies examining the relative benefits of sutures, staples, adhesive strips, and glues for stability, infection, cosmesis, and other factors [52–57, 146–151], but there has been little consideration of pain-outcomes [54, 146]. Pain was considered in the context of closure of oral wounds [151]. We have found that glue-closure obviates the nerve-injury-like changes in gene expression that are associated with suture- and staple-closure of skin incisions. The suite of data we have produced around the effects of skin incision on sensory neurons suggest that different closure methods should be considered as a factor in long-term pain outcomes.

### Considerations for PEG Effects

#### Does PEG act to prevent or reverse expression of ATF3?

We did not design an experiment to address this question. However, there are indications from separate experiments that can address this important question. The result for PEG-treatment in terms of ATF3 mRNA expression appears inconsistent with the results for neurons expressing ATF3 protein in the PEG-treated animals. One possibility is that the protein could have been expressed due to some injury which occurred prior to PEG treatment, and PEG did not reverse that expression. Our prior work demonstrated that a small proportion of DRG neurons are the injured by the process of injecting the DiI tracer into the skin, and they express ATF3 protein even without incision [10]. Indeed, the proportion of ATF3+ neurons remaining after PEG-treatment (**Figure 12C**) is similar to naive with DiI-tracing. This appears to be the most-likely scenario when the incision is separated in time from the DiI injection. It would imply that PEG-treatment is acting to prevent induction, and not reversing the effect. If the incision were not significantly separated in time, the ATF3+ neurons might still have been induced prior to the PEG-treatment, but this is less likely. Thus, these combined data suggest that PEG treatment (as performed here) more likely acts to prevent induction of ATF3 than reversing/suppressing its ongoing expression.

Taken together, PEG-treatment dramatically reduces the incision-induced and pain-associated molecular and functional indications of axonal injury and cellular stress. ATF3 mRNA from bulk tissue is reduced, as is the number of incision-associated neurons expressing ATF3 protein. Although the electrophysiological sensitization of responsiveness of the individual ATF3+ neurons is still notable, *there are far fewer of those neurons remaining after PEG treatment*. Further, the PEG-treatment at the time of the incision (28 days earlier) stabilized the threshold for activation (rheobase), suggesting a prevention of the hypersensitivity that usually occurs after incision (for neurons expressing ATF3), as opposed to a delay.

### Treatment-Mediated Reduction in ATF3

The number of ATF3+ sensory neurons 3 weeks after induction of osteoarthritis in two separate models was reduced by treatment with a LPA receptor inhibitor [152]. Since inhibitor dosing included both pre- and post-OA-induction treatment, the effect on ATF3 expression was likely via prevention, and not reversal. Another study [153] determined that delayed administration of TLR4-antagonist could reduce the number of ATF3+ neurons in the relevant DRGs. It was unclear if this was due to prevention of de novo induction of ATF3 in a new population of neurons in the tissue degeneration model or a reversal of expression already induced. Nonetheless, it is compelling that other treatments are also capable of reducing the overall expression of ATF3 in sensory neurons.

### Study Limits

Although we modeled the ketoprofen regimen from clinical use and prior experimental protocols, we did not directly characterize the nature or degree of inflammation present in this model. While we can be very confident that the ketoprofen treatment had anti-inflammatory effects, we cannot be certain of any specifics about the effects of that treatment. Although this is not ideal, the conclusion that inflammation is neither necessary not sufficient is not particularly weakened. If inflammation were sufficient and the ketoprofen treatment had no effect, then we would expect to see increased ATF3 expression induced by incisions close-outside of the dermatomal border (inflammation without axon injury), but this did not occur (**Figure 2**).

It must be noted that the model used here is one with mixed tissue – the incision injures both the skin and the attached underlying CTM. The exact impact of this is not known, but it is worth mentioning as Brennan and colleagues have reported that incision-related pain is more severe when the damage is to both skin and muscle, as opposed to only skin [27]. However, they also note that “…a skin incision without a muscle tissue injury seems to be responsible for inducing mechanical hyperalgesia after incision; muscle injury seems to be not required”[6]. One would expect that, if inflammation alone were capable of inducing ATF3 expression, it would be more likely in the case of incision of both skin and muscle as opposed to skin alone [47, 134], further reinforcing our conclusion that inflammation is not necessary for ATF3 induction. In terms of clinical relevance, cutaneous muscle coverage is highly variable in humans and is not something that is routinely considered when planning and executing most surgeries, or in follow-up for post-surgical pain.

Although there has been extensive work done to understand the ways that PEG may influence biological systems [20, 21, 64, 65, 67, 154–162], the mechanisms underlying the PEG effects here are unknown. The experimental design was decidedly broad and not refined. Given that a known anti-inflammatory agent did not prevent ATF3-induction, but PEG did prevent ATF3 induction, we presume that PEG is acting through some mechanism other than preventing inflammation (which was shown to be insufficient for inducing ATF3 expression anyway). The presumption, and the rationale underlying the initial design, is that PEG is acting to enhance membrane integrity and perhaps seal the axolemma injured by the incision. If this were actually occurring it could perhaps prevent or delay some positive or negative injury signal from causing changes in gene expression at the DRG. In essence, it may be “fooling the neuron into thinking it isn’t injured”.

Encouragingly, PEG-treatment also led to a reduction of incision-associated electrophysiological changes, but it is not clear if this is due to prevention of ATF3-induction (and thus to the presumed-but-not-proven effects on gene expression), to direct effects on the membrane properties, or both. Importantly, we also examined only a limited time window. PEG effects on ATF3 expression were examined at 4d (mRNA) and 28d (protein), and electrophysiological properties only at 28d. The reduction of ATF3 at both 4d and 28d is compelling and suggests that the effect of PEG is to prevent ATF3 induction. It is nonetheless possible that PEG treatment may just delay ATF3 expression and emergence of electrophysiological sensitization or both, and not fully-prevent it. This will require additional assessments.

We have not yet examined whether autonomic neurons and motor neurons are similarly affected. The focus on sensory neurons is nonetheless highly legitimate given the topic of pain. However, future examination of the other PNS populations is warranted considering the known crossed-influence that can occur, particularly in conditions in which tissue micro-scale compartments can be compromised such as incision and neuroma.

As an aside, it is possible that PEG-treatment might be a useful way of extending the “acutely-dissociated” status of adult sensory neurons, which start to display injury-associated transcriptional and electrophysiological changes after 8-12 hours in vitro (varying with species and culture temperature) [76].

Although we did not observe ATF3 protein expressed in non-neuronal cells, it is certainly possible that this nonetheless occurred either below visual detection threshold or it was expressed at a time we did not examine. Recent work has tied ATF3 expression by macrophages to persistent neuropathic pain [163]. Macrophages infiltrate the DRG and localize near injured sensory neurons after overt nerve injury, but it is not clear if this occurs with incision or other forms of damage to peripheral target tissues. Clarifying this unknown would be very important for determining the relationship of the sensory system response to tissue damage to that of nerve injury.

### Translating animal models to clinical outcomes

”A different paradigm is required for the identification of relevant targets and candidate molecules whereby pain is coupled to the cause of sensorial signal processing dysfunction rather than clinical symptoms” “Given that neuropathic and chronic pain results from a preceding dysfunction in sensory signalling, the identification of effective treatments requires further insight into the reversibility of the underlying dysfunction as well as the timing of intervention relative to the onset of the disease. Novel therapeutic interventions need to be focused at the dysfunction in signalling pathways rather than primarily on pain relief.”[164]

The general approach of the research – and even moreso the clinical – fields is that the various forms and players of inflammation are the root of the emergence of persistent pain (e.g., [1, 2]). Inflammatory and neuropathic pain are rightfully considered different because they result from distinct mechanisms. Inflammatory pain is generally associated with injury to peripheral tissues and neuropathic pain generally associated with injury to the brain, spinal cord, and nerve tissues. The data presented here, combined with those from others, suggest that damage of peripheral tissues might be inducing biological responses and/or conditions that could lead to neuropathic pain, with or without inflammation and inflammatory pain. We do not debate a role for tissue or neuro-inflammation in acute pain or persistent pain. New categories of pain with terms have been proposed, including “nociplastic”. We are not advocating for the biological responses here to be considered as mechanisms of “nociplastic pain” (e.g., [165]), though there are features of “nociplastic pain” that make that possibility both feasible and attractive. Instead, we view these and our prior data [9, 10] as supporting a revised consideration of the impact of tissue damage on the nervous system. This revised consideration could aid in generating a better mechanistic classification of persistent pain after tissue damage. We also lean on many in the field who recognize that there is insufficient mechanistic understanding for how inflammatory processes alone lead to observed phenotypes (e.g., [6, 78]). In accord with recommendations for using mechanism-based approaches to pain diagnosis and treatment (e.g., [166]), we propose that refining the concept, and recognizing additional biological processes such as we describe here, may better account for the observations and provide a better framework for applying basic research to clinical needs as regards persistent pain after tissue damage and surgery (**Figure 14**). More work needs to be done to determine if there are meaningful differences in the specific responses to tissue injury-induced axonal damage and nerve injury-induced axonal damage, but the conceptual refinement of considering the long-term consequences of tissue damage as more-similar to nerve injury than to inflammation may provide a significant advance.

**Figure 14.**
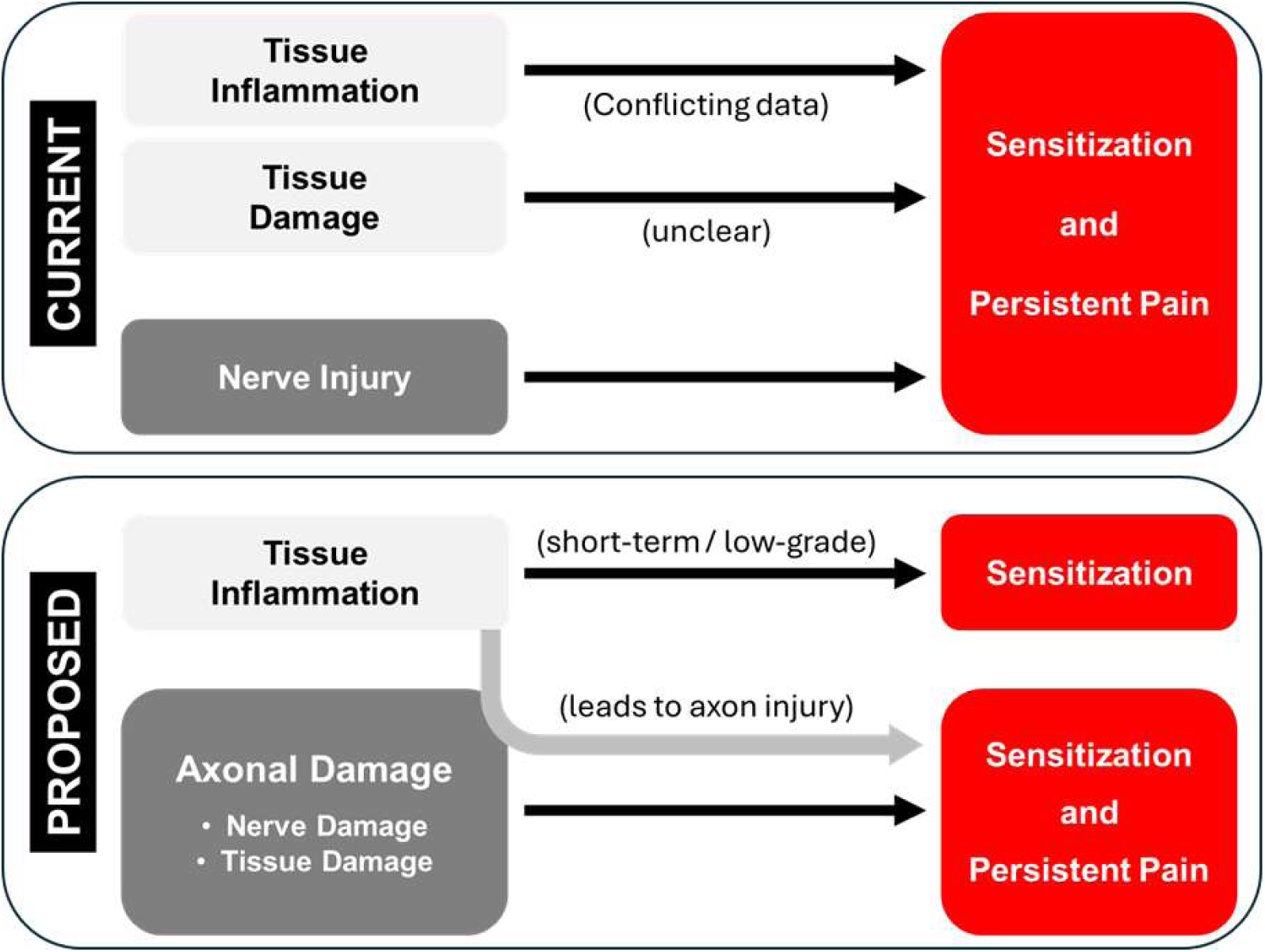
Schematic of the general concept of the relationship of inflammatory and injury conditions to the emergence of sensitization and persistent pain. Current concept is contrasted with the proposed conceptual change. Most notable is inflammation alone would be insufficient to lead to persistent neuropathic pain without concomitant axonal damage, which would unify how nerve injury and tissue damage can lead to persistent neuropathic pain even after wounds are healed and inflammation is resolved or suppressed. This model does not incorporate the many known analgesic/anti-nociceptive mechanisms.

We also used the experimental model and concepts espoused here to provide preliminary indications that simple, feasible, and expandable approaches might be used to prevent some of the biological responses that could be contributing to persistent pain.

### Future Directions

Targeting transcription factors can be difficult, but this approach for pain control directed to the DRG and spinal dorsal horn already has proof of concept [167]. It is possible that directed interference with ATF3 function in sensory neurons may be an effective approach for preventing and/or treating persistent pain after tissue damage. Assessing the feasibility of this approach will require defining the temporal profile of ATF3 and other axon injury-associated transcriptional regulators and determine how they interact to control transcription leading to intrinsic sensitization. There is proof of concept for prevention of ATF3 induction in models of osteoarthritis by pre-treatment with a signaling inhibitor [152] or reduced increase by later administration of an inhibitor of TLR4 [153].

Encouragingly, closing wounds with glue – a common option for many surgeries and injuries – appears to reduce markers of lost-lasting sensory neuron sensitization. This may reduce the risk of developing persistent pain. Further advances in tissue-adhesion / wound-repair may affect the responses reported here. Treating tissue damage with agents like PEG could also become part of the pain-prevention regimen with surgery or trauma [17, 168–174]. Although we tested PEG effects with both systemic and topical administration, it is entirely feasible that topical application alone might have similar/suitable effects. Assessing the feasibility of this approach will require directly testing different routes of administration and determining the temporal therapeutic window.

## Acknowledgements

Renee Donohue

KSCIRC Cores

Jon D. Levine for encouragement at early stages

